# A Developmental Lectin-Glycan Program Enables Early Breast Cancer Dissemination and Metastatic Onset

**DOI:** 10.64898/2026.02.23.707584

**Authors:** Ramiro M. Perrotta, Magalí Berton, Luis E. Valencia Salazar, Yamil D. Mahmoud, Juan M. Pérez Sáez, Tomás Dalotto-Moreno, Beatriz Del Valle-Pérez, Neus Martínez-Bosch, Rosa M. Morales, Sabrina G. Gatto, Pilar Navarro, Raquel Batlle, Julio A. Aguirre-Ghiso, Gabriel A. Rabinovich, Mariana Salatino

## Abstract

Early dissemination of breast cancer cells can precede clinically detectable tumor progression, yet the programs enabling this process, remain poorly understood. Here, we identify a developmental glycocheckpoint governed by the galectin-1 (GAL1)-glycan axis that is hijacked during early breast cancer dissemination. In the mammary gland, regulated GAL1 expression and glycan accessibility directed epithelial lineage specification and progesterone-induced branching morphogenesis. This program was aberrantly reactivated in early breast cancer lesions to promote epithelial plasticity, stem-like traits and metastatic competence. Mechanistically, GAL1 was enriched in mammary stem cell compartments and sustained progesterone receptor expression and activity. Genetic ablation or therapeutic inhibition of GAL1 across breast cancer models restrained early lesion progression, reduced circulating tumor cell frequency, and limited lung metastasis. Consistent with these findings, high GAL1 expression combined with low expression of the GAL1-restricting sialyltransferase ST6GAL1 was associated with poor clinical outcomes in patients. Thus, breast cancer co-opts a developmental GAL1-glycan program to disseminate early, revealing an unexpected link between mammary morphogenesis and metastatic progression and identifying GAL1 as a therapeutic vulnerability in early-stage disease.

## INTRODUCTION

Despite substantial progress in systemic therapies, metastatic breast cancer remains largely incurable. Disease relapse and therapeutic failure frequently occur even after initial clinical responses, suggesting that key biological programs enabling dissemination and metastatic competence are established prior to treatment and before clinically detectable metastases emerge ^1,2^. These limitations highlight the urgent need to identify early molecular drivers of dissemination that could be targeted to intercept metastatic progression.

During normal tissue development, epithelial cells engage highly coordinated morphogenetic programs that regulate lineage specification, collective migration, and tissue architecture. In the mammary gland, these processes drive ductal elongation and branching morphogenesis through dynamic regulation of epithelial plasticity, cell-cell interactions, and extracellular matrix remodeling ^3,4^. Notably, mammary branching morphogenesis shares striking phenotypic parallels with processes associated with cancer cell invasion and dissemination, including coordinated migration, transient disruption of epithelial organization, and reversible changes in cell identity ^5^. These processes have been largely studied at the genetic and epigenetic levels ^6,7^.

Glycosylation, the process by which glycans are covalently attached to proteins and lipids, plays a critical role in regulating cell signaling, adhesion, and immune recognition ^8^. Aberrant glycosylation is a critical determinant of cancer and contributes to different hallmarks including invasion, angiogenesis, metastasis, and immune evasion ^9^. Galectins, a family of β-galactoside-binding lectins, have emerged as key decoders of glycan information and potent modulators of the tumor microenvironment (TME) ^10^. Developmentally regulated and elevated at sites of tumor growth and metastasis, galectins operate through both intracellular and extracellular mechanisms to foster progression of the metastatic cascade ^11^. Galectin-1 (GAL1; encoded by the *LGALS1* gene), a prototype member of the galectin family, binds to terminal N-acetyllactosamine (LacNAc) residues present in both complex N- and O-glycans ^12^. This glycan-binding protein influences several processes such as epithelial-to-mesenchymal transition (EMT), pathological angiogenesis and immunosuppression ^12–19^. GAL1 overexpression has been associated with resistance to chemotherapy, radiotherapy, anti-angiogenic therapies and immunotherapies, highlighting its relevance as a broad therapeutic target in cancer ^20^.

The accessibility to GAL1 binding is governed by the concerted action of glycosyltransferases, whose expression and activity are differentially regulated in the tumor microenvironment (TME) ^21^. Whereas mannoside acetylglucosaminyltransferases (MGATs) generate N-glycan branching, β-1,4-galactosyltransferase (B4GALT) and β-1,3-N-acetylglucosaminyltransferase (B3GNT) elongate LacNAc repeats, and sialyltransferases (ST) promote sialylation. Specifically, incorporation of terminal α2,6-linked sialic acid by the β-galactoside α-2,6-sialyltransferase 1 (ST6GAL1) inhibits GAL1 recognition of LacNAc motifs ^12^. While elevated ST6GAL1 expression has been linked to tumor progression in several contexts, recent evidence has shown that its loss in circulating tumor cells (CTC) enhances chemo-evasion, clustering, and metastatic seeding in triple-negative breast cancer, underscoring the context-dependent complexity of this glycosylation-driven circuit ^21,22^.

Here, we show that a progesterone-driven GAL1-glycan signaling axis governs mammary branching morphogenesis, a key process in mammary gland biology, and is co-opted by early breast cancer lesions to promote dissemination and metastatic competence. Remarkably, we further uncover a non-canonical nuclear function of GAL1 as a direct regulator of PR expression, revealing a connection between lectin biology and hormonal control of mammary development. Analysis of *MMTV-PyMT* and *MMTV-Neu* transgenic mouse models together with patient-derived clinical data, revealed a critical role of GAL1 in promoting epithelial plasticity, stem-like traits, and early dissemination, uncovering a mechanistic link between developmental morphogenesis and early metastatic dissemination. Systemic administration of a neutralizing anti-GAL1 monoclonal antibody (mAb) lowered circulating GAL1 levels, impaired mammary branching, decreased CTC burden, and suppressed lung metastasis.

Together, these findings identify GAL1 as a regulator of mammary gland cell homeostasis and morphogenesis that is usurped by tumors to drive dissemination. Furthermore, we propose that a *LGALS1/ST6GAL1* signature might serve as a candidate biomarker and a druggable axis for antimetastatic interventions in breast cancer.

## RESULTS

### Endogenous GAL1 regulates progesterone-dependent mammary gland branching and morphogenesis

Mammary gland branching morphogenesis is a complex developmental process where normal epithelium proliferates and invades the surrounding stroma through terminal end buds (TEBs) into an extensive tree-like ductal network ^23^. Estrogens and progesterone coordinate this invasive process and can be co-opted by tumor cells to disseminate and seed metastases ^1,2^. GAL1 has been implicated in normal branching ^24^ and in cancer cell migration and invasiveness ^11^. To interrogate whether tumor cells may hijack the GAL1-glycan pathway for dissemination, we examined its shared mechanistically linked roles in normal gland development and early tumor progression.

We first compared ductal branching in GAL1-deficient (*Lgals1*^-/-^) and wild-type (*Lgals1^+/+^*) mice (*C57BL/6J*) following administration of the progesterone analog medroxyprogesterone acetate (MPA). *Lgals1^-/-^* mice displayed significantly reduced branching at baseline and after MPA, characterized by a lower number of TEB (Figure 1A, 1B) and a lower branching index (Figure 1A, 1C).

**Figure 1:**
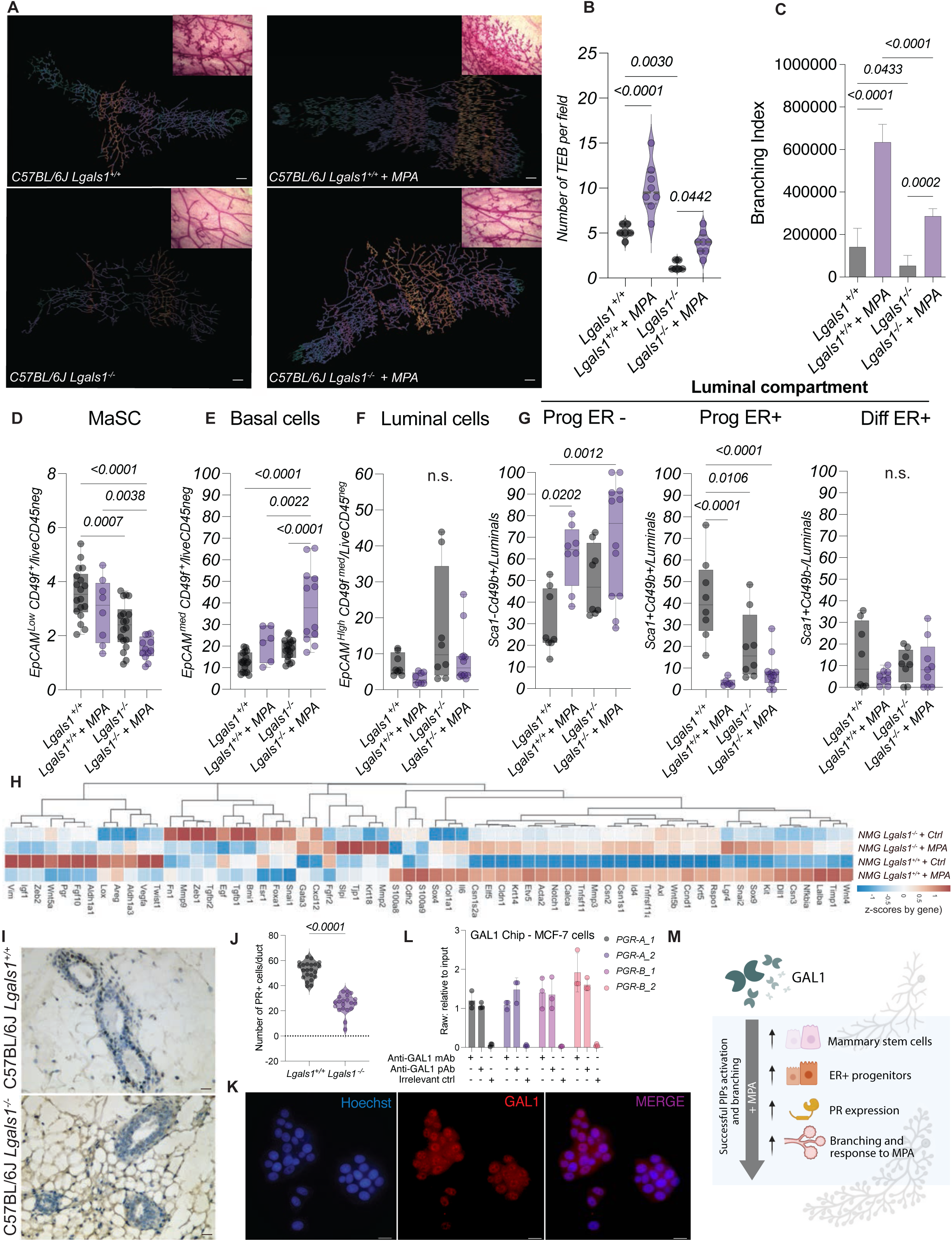
**GAL1 remodels cell composition and promotes morphogenesis of normal mammary gland**. A) Representative whole mount of mammary glands (MG) from C57BL/6J *Lgals1*^+/+^ or *Lgals1*^-/-^ mice treated or not with MPA. Images were generated using Sholl mask tool. Scale bar, 2 mm. B) Quantification of MG branching as the number of terminal end-buds (TEB) per field. C) Branching index. D-F) Flow cytometric analysis of epithelial lineage markers in *Lgals1*^+/+^ and *Lgals1*^-/-^ MG, showing the percentage of: D) mammary stem cells (MaSC, EpCAM^low^ CD49f^+^), E) basal cells (EpCAM^med^ CD49f^+^), and F) luminal cells (EpCAM^high^ CD49f^low^), with or without MPA stimulation. G) Analysis of the MG luminal compartment (ER^+/-^ luminal progenitors and differentiated ER^+^ cells) based on Sca1 and CD49b expression, H) Heatmap displaying mRNA expression *z* scores of MG morphogenesis includingrelevant genes measured by RNAseq. I-J) Representative images of PR staining by immunohistochemistry (IHC) Scale bar, 20 μm. (I) and determination of PR^+^ cells per duct (J). K) Representative immunofluorescence images showing GAL1 expression (red) in MCF7 breast cancer cells. Nuclei were counterstained with Hoechst (blue). Scale bar, 20 μm. L) ChIP-qPCR validation of GAL1 enrichment at progesterone receptor (*PGR*) promoter regions. Four primer pairs spanning *PGR* regulatory regions (two targeting *PGRA* and two targeting *PGRB*) were used to assess enrichment (mAb: monoclonal and pAb: polyclonal, anti-GAL1 antibodies). M) Schematic representation of GAL1 effects on MG branching morphogenesis. **Statistical analysis:** Data are presented as mean ± SD. For panels B, C, E, F, G and L, differences between groups were analyzed by one-way ANOVA followed by Tukey’s post-test. (H) n = 3 mice per group. (J) For panel J, an unpaired Welch’s t-test was used to compare *Lgals1^+/+^* and *Lgals1^-/-^* MG. p-values < 0.05 were considered statistically significant. All experiments were performed with n ≥ 5 mice per group.

Normal ducts contain two epithelial cell types: hormone-responsive luminal cells lining the lumen and basal myoepithelial cells contacting the basement membrane ^23^. As the TEB advances, mammary stem cells (MaSC) contribute to ductal expansion by leaving progeny cells behind ^25^. To assess whether endogenous GAL1 alters epithelial cell hierarchies, we profiled epithelial cell populations by flow cytometry using EpCAM, CD49f, and Sca1 ^26,27^ in *Lgals1^-/-^* and *Lgals1^+/+^* animals (Figure S1A). These phenotypic markers resolved four central compartments: basal (EpCAM^med^ CD49f^+^), luminal (EpCAM^+^ CD49f^low^), MaSC-enriched (EpCAM^low^ CD49f^+^) and stromal (EpCAM^-^ CD49f^-^) (Figures 1D, 1E, 1F, S1B). Lineage identities were confirmed by RT-PCR in sorted cells: luminal (*Progesterone Receptor; PR*), basal (*Acta2*), and MaSC (*Nanog*) (Figure S1C). GAL1 expression was enriched in the basal and MaSC populations indicating that these compartments may serve as major sources of this lectin in the gland (Figure S1D). The absence of GAL1 disrupted this epithelial cell hierarchy in the mammary gland as *Lgals1^⁻^*^-/-^ mice exhibited a reduced MaSC population. This effect was further pronounced after MPA treatment, consistent with impaired stem cell expansion or maintenance (Figure 1D) and the observed branching deficit (Figure 1A, 1B and 1C). In contrast, MPA treatment led to an expansion of the basal cell compartment in *Lgals1*^-/-^ glands, pointing to altered lineage dynamics and hormone responsiveness (Figure 1E).

Although no significant differences were detected in the overall luminal compartment (Figure 1F), stratification of luminal subtypes using Sca1 (Ly6a) and CD49b as surrogate markers of ER^+^ and ER^-^ progenitor cells, ^27,28^ revealed distinct alterations between genotypes. *Lgals1^-/-^* mice showed a decreased proportion of ER^+^ luminal progenitors (Sca1⁺CD49b⁺), potentially compromising paracrine cues from hormone-sensing cells (Figure 1G). As expected, MPA expanded ER-luminal progenitor cells (Sca1⁻CD49b⁺) in both genotypes. Fully differentiated non-clonogenic ER^+^ luminal cells (Sca1^+^CD49b^−^) were not affected by the absence of GAL1 (Figure 1G). These findings suggested a possible role for GAL1 in proper progesterone-driven remodeling of the mammary epithelial hierarchy and branching morphogenesis.

RNA sequencing (RNAseq) analysis revealed that GAL1 regulates progesterone-induced transcriptional remodeling during mammary gland development (Figure 1H, Table S1). In response to MPA treatment, *Lgals1*^+/+^ mammary tissues underwent extensive transcriptional reprogramming, with the induction of genes involved in alveologenesis, milk protein synthesis, immune modulation, and epithelial cell proliferation. Canonical progesterone-responsive genes, including *Tnfsf11* (*Rankl*) and *Wnt4*, *Elf5* and *Calca* and casein-encoding genes (*Csn2, Csn1s1, and Csn1s2a*) ^29^ were strongly upregulated by MPA in *Lgals1*^+/+^ mice (Figure 1H, Figure S1F). In contrast, GAL1-deficient glands exhibited attenuated responses, with reduced induction of *Rankl*, *Elf5*, and *Rspo1*, genes that are critical for MaSC renewal and alveolar differentiation ^30–32^ (Figure 1H, Figure S1E). This attenuated transcriptional response is specific to the progesterone-induced condition and does not reflect a global difference in gene expression, since baseline gene expression in vehicle-treated tissues was comparable between genotypes (Figure 1H).

Surprisingly, we detected a marked reduction in PR expression in the mammary gland of *Lgals1^-/-^* mice by immunohistochemistry (IHC) (Figures 1I, J), RNAseq (Figure S1E) and RT-PCR (Figure S1F), providing a mechanistic link to the impaired response to MPA-induced branching and MaSC expansion. To investigate possible mechanisms linking GAL1 to PR expression, we examined whether GAL1 associates with the *PGR* promoter A and B in human breast cancer cell lines. Subcellular fractionation revealed the presence of GAL1 in the cytoplasm, as expected, but also within the nucleus and the chromatin fraction of both PR^+^ MCF7 (Figure 1K, Figure S1G) and T47D cells (Figure S1H, S1I). Consistently, ChIP-qPCR assays performed with two independent anti-GAL1 antibodies, demonstrated significant enrichment of GAL1 at both *PGR* promoters A and B, compared with an irrelevant IgG control in MCF-7 (Figure 1L) and T47D cells (Figure S1J). These findings are compatible with a model in which chromatin-associated GAL1 may participate in the transcriptional regulation of *PGR*, providing a potential mechanistic explanation for the reduced PR expression observed in *Lgals1*-deficient mammary glands.

These findings suggest that GAL1 is a critical regulator of hormone-driven mammary gland development, capable of supporting PR expression, MaSC reservoir, and the progesterone-responsive transcriptional program that underlies branching morphogenesis (Figure 1M). Expression of *Rankl*, *Csn2*, and *Csn1s2a* in *Lgals1^-/-^* tissues under basal conditions, indicates that GAL1 is critical for transcriptional responses to hormonal cues, particularly those delivered by progesterone. By enforcing this physiologic checkpoint, GAL1 may provide a developmentally regulated pathway that tumor cells may later co-opt to enable early dissemination. To investigate the spatiotemporal dynamics of GAL1 in the mouse MG, we interrogated single-cell RNA sequencing (scRNA-seq) datasets spanning embryonic development throughout adulthood ^33^ (Figure 2A). Consistent with our IHC/flow cytometry data in adult MG (Figure S1E), *Lgals1* transcripts were enriched in the basal and MaSC lineages (Figure 2A, 2B and 2C) and positively correlated with the basal marker *Acta2* (α-SMA) (Figure 2B; Figure S2A).

**Figure 2:**
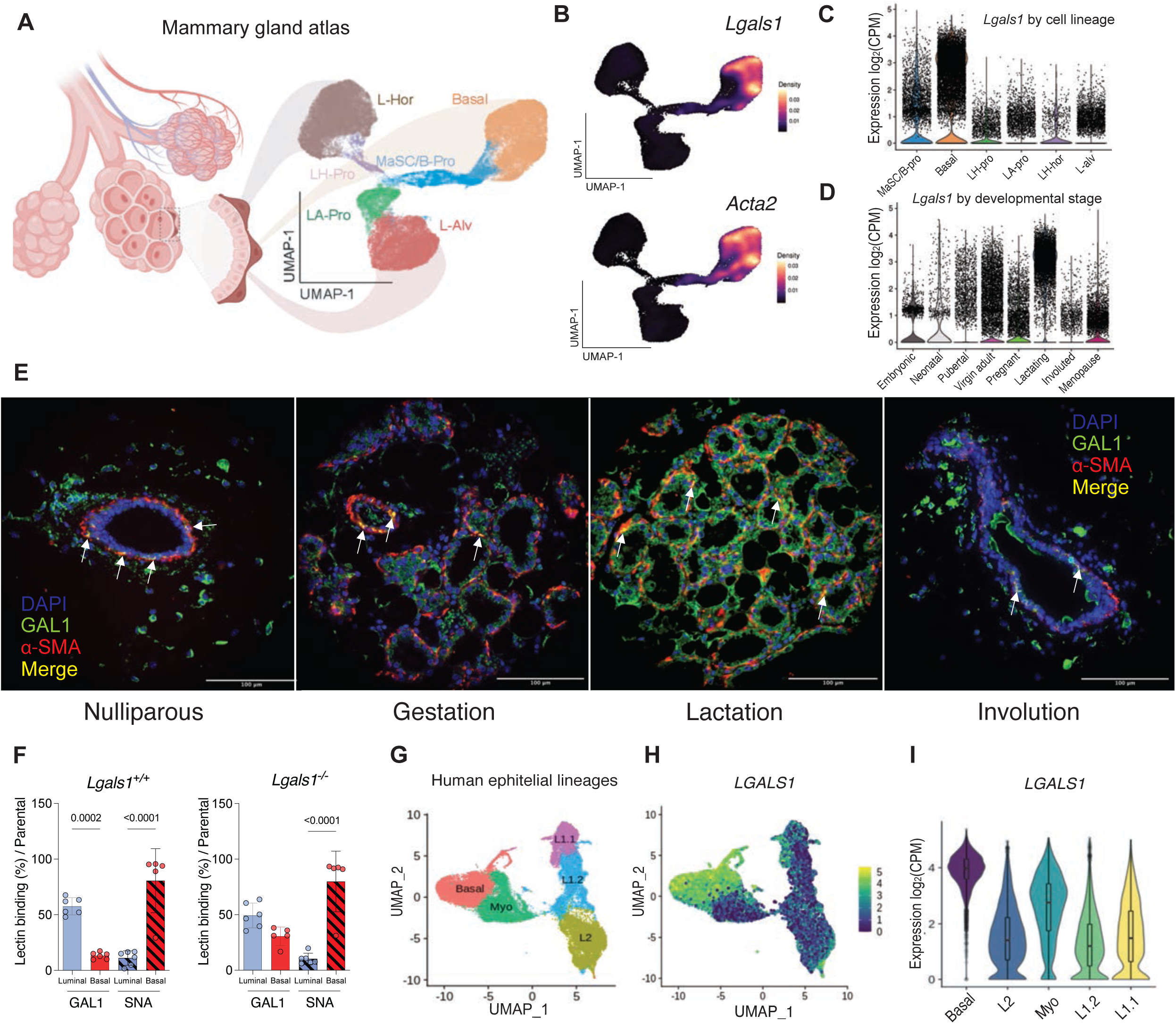
Transcriptomic analysis of MG reveals differential GAL1 expression across different epithelial cell lineages and stages. A-B) A 2D Uniform Manifold Approximation and Projection (UMAP) of the ATLAS data is organized into clusters, classified according to murine mammary cell lineages. L-Hor: luminal hormone-sensing cells; L-Alv: luminal alveolar cells; Basal: basal cells; LH-Pro: luminal hormonal progenitor; LA-Pro: luminal alveolar progenitor; MaSC/B-Pro: mammary stem cells / basal progenitors. B) Expression of *Lgals1* and *Acta2* (α-SMA) is visualized on the UMAP projection. C-D) *Lgals1* expression across the different cell lineages (C) or developmental stages (D) identified in the MG. E) Representative immunofluorescence images showing expression of α-SMA and GAL1 in MG of C57BL/6J mice at different developmental stages. DAPI was used for nuclei staining. F) Flow cytometry analysis of GAL1 and SNA binding in C57BL/6J *Lgals1^+/+^* or *Lgals1^-/-^* mice MG. G) UMAP projection color-coded by clusters corresponding to human mammary cell lineages. The UMAP was generated using 15 dimensions and a cluster resolution of 0.01 with Seurat V4. H-I) Expression of *LGALS1* across the previously defined cell lineages.

Across developmental stages*, Lgals1* showed sharp regulation, higher relative expression during embryonic and neonatal stages, declined at puberty, and increased again in adulthood, with a second peak during lactation (Fig 2D, Figure S2B). This dynamic regulation suggests a potential role for GAL1 in both ductal morphogenesis and alveolar maturation. Confocal microscopy of normal C57BL/6J MG at different physiological stages (nulliparous, gestation, lactation, involution) confirmed basal layer localization of GAL1 together with α-SMA (Figure 2E).

RNA velocity analysis ^34^ revealed that GAL1 is dynamically expressed and transcriptionally active within basal and MaSC populations reaching its peak expression in the basal cell lineage, consistent with our phenotypic characterization (Figure S2C and S2D). In contrast, ST6GAL1, an enzyme that catalyzes the α2,6-sialylation of terminal LacNAc residues, thereby masking GAL1-binding sites, showed reduced expression in the hormone-responsive luminal (L-Hor) population, with increased expression toward the basal lineage (Figure S2E). Consistently, glycophenotypic analysis revealed lower binding of the *Sambuccus nigra* agglutinin (SNA) in the luminal lineage; further indicating the reduced frequency of α2,6-linked sialic acid on these cells. Consistent with this sialophenotype, GAL1 binding was higher in luminal cells (Figure 2F). Pseudotime and dynamical modeling further identified *Lgals1* mRNA as one of the top enriched transcripts within the basal lineage during periods of active morphogenesis (Figure S2F-G; Table S2). Analysis of additional scRNA-seq datasets ^35^, revealed that mammary stromal populations, including fibroblasts and immune cells such as macrophages, NK cells, and conventional dendritic cells, also represent significant sources of GAL1 (Figure S2H), suggesting a broader role for this lectin within the MG microenvironment. To elucidate the relevance of GAL1 in the human MG, we then analyzed the spatiotemporal expression of this lectin using a publicly available scRNAseq dataset of normal human MG ^36^. Consistent with our findings in mouse settings, this atlas resolves basal cells and three luminal subsets: luminal progenitors (L1.1), mature secretory alveolar cells (L1.2), and hormone-responsive luminal cells (L2), defined by EpCAM and CDH1 phenotypic markers (Figure 2G). Notably, *LGALS1* exhibited the highest expression in basal and myoepithelial cell populations, mirroring the distribution observed in the mouse MG (Figure 2H and 2I).

Together, these data provide transcriptomic and spatiotemporal evidence suggesting that GAL1 contributes to basal and MaSC identity, being dynamically regulated across mammary development. This spatiotemporal expression pattern supports an essential role for GAL1 in hormone-driven remodeling and branching morphogenesis of the MG, with potential implications for breast cancer initiation and early progression.

### The *LGALS1/ST6GAL1* expression ratio identifies a high-risk group in human breast cancer

Building on our findings that GAL1 regulates MG morphogenesis and hormone-driven branching and given that developmental programs are frequently co-opted during tumorigenesis ^36^, we next investigated whether GAL1 contributes to breast cancer progression in patients.

To assess the prognostic value of GAL1, we first examined disease-free survival as a function of *LGALS1* expression in a cohort of patients. Survival analyses were conducted using the TCGABRCA cohort (n = 977), which includes primary breast tumors with matched gene expression and clinical outcome data (STAR Methods, Table S4). Stratification into *LGALS1*^high^ and *LGALS1*^low^ groups (median *LGALS1* expression; Table S4) did not show an association with clinical outcome, indicating that *LGALS1* expression alone is insufficient to stratify patient risk (Figure 3A). Given that GAL1 binding to cell surface glycans is functionally constrained by α2,6-sialylation ^12^, we next evaluated the impact of *ST6GAL1* expression on patient survival. *ST6GAL1* expression alone demonstrated a significant association with improved survival, with high *ST6GAL1* expression conferring an increased DFS (Mantel-Cox p = 0.0189; hazard ratio [HR] = 1.57) (Figure S3A).

**Figure 3:**
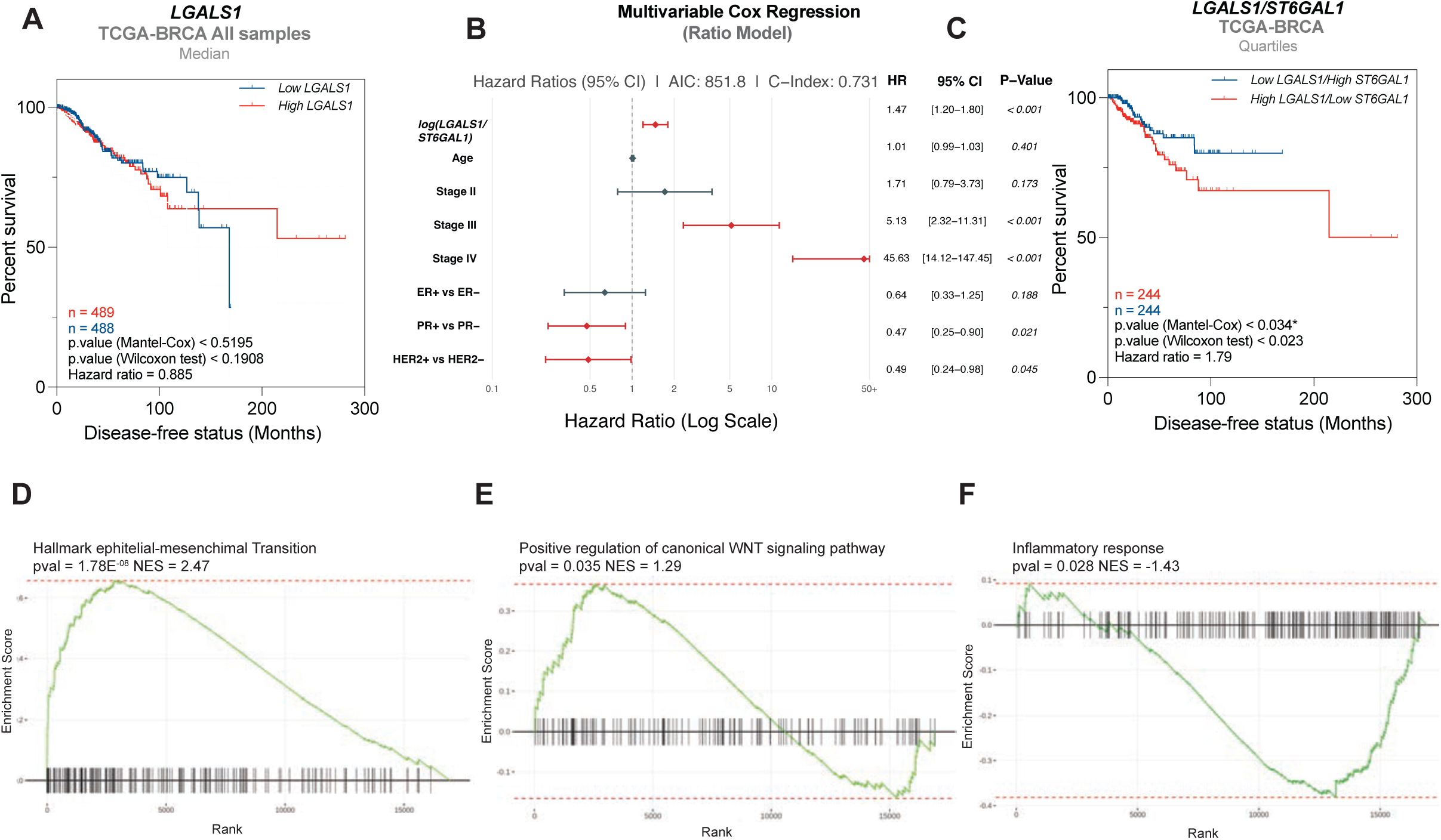
***LGALS1/ST6GAL1* ratio is associated with clinical outcome in breast cancer.** A) Kaplan-Meier DFS survival curve of patients with *LGALS1^high^* and *LGALS1^low^*expression (stratification: median *LGALS1* expression). B) Multivariate Cox proportional hazards regression analysis of the log-transformed *LGALS1/ST6GAL1* ratio in the TCGA-BRCA cohort. Forest plot showing hazard ratios (HR) and 95% confidence intervals for disease-free survival, adjusted for age, tumor stage, and receptor status (ER, PR, HER2). The vertical dashed line indicates HR = 1 (no effect). Variables with HR > 1 are associated with increased hazard, whereas HR < 1 indicates a protective effect. C) Kaplan-Meier DFS survival curve of patients with *LGALS1^high^/ST6GAL1^low^*and *LGALS1^low^/ST6GAL1^high^* expression. D-F) Enrichment plots of: D) the epithelial-to-mesenchymal transition (EMT) hallmark (GSEA); E) positive regulation of Wnt canonical pathway and F) inflammatory response hallmark. Data is presented as biological processes up- or down-regulated in the *LGALS1^high^/ST6GAL1^low^* group. **Statistical analysis:** (A and C) Disease-free survival was compared using the Mantel-Cox log-rank and Wilcoxon tests; hazard ratio (HR) was calculated from the survival curves. (B) Statistical significance was set at p < 0.05. All hazard ratios (HR) are reported with 95% confidence intervals (CI). (D-F) Pathways significantly modulated in the GSEA with P < 0.05 were considered significant.

To determine whether the balance between *LGALS1* and *ST6GAL1* expression independently associates with clinical outcome, we next performed multivariate Cox proportional hazards regression incorporating age, tumor stage, and receptor status (ER, PR, HER2). Modeling the log-transformed *LGALS1/ST6GAL1* ratio as a continuous variable revealed that a higher ratio was significantly associated with worse disease-free survival ([HR] = 1.47, 95% CI 1.20-1.80, p < 0.001), independent of established clinicopathologic covariates (Figure 3B). In contrast, when the genes were modeled individually, *ST6GAL1* expression retained an independent protective effect (HR = 0.63, 95% CI 0.46-0.86, p = 0.003), whereas *LGALS1* showed only a trend toward association (HR = 1.36, 95% CI 0.98-1.89, p = 0.062) (Figure S3B). While both models offered similar discrimination (identical C-index), the ratio-based model provided a superior statistical fit (lower Akaike Information Criterion [AIC], where lower values indicate better model fit) and stronger statistical evidence (lower p-value), indicating that the interplay between *LGALS1* and *ST6GAL1* better explains survival variability than modeling each gene independently.

Comparison of the biologically opposed extreme states, *LGALS1*^high^/*ST6GAL1*^low^ versus *LGALS1*^low^/*ST6GAL1*^high^, revealed a more pronounced segregation in survival than that observed for expression of individual genes alone, consistent with a context-dependent interaction between *LGALS1* expression and α2,6-sialylation status. Specifically, the *LGALS1*^high^/*ST6GAL1*^low^ group exhibited significantly increased hazard relative to the *LGALS1*^low^/*ST6GAL1*^high^ group (Mantel-Cox p = 0.034; HR = 1.79) (Figure 3C, Figure S3B), indicating the worst clinical outcome.

Together, these data support a context-dependent model in which *LGALS1* acquires prognostic relevance specifically in tumor microenvironments (TME) with reduced *ST6GAL1* expression. Thus, GAL1 expression becomes clinically meaningful when α2,6-sialylation is mitigated. To understand the molecular programs enriched in this high-risk group, we performed Gene Set Enrichment Analysis (GSEA). Patients with *LGALS1^high^/ST6GAL1^low^*phenotype showed a significant enrichment for Epithelial-to-Mesenchymal transition (EMT) programs (Figure 3D left), and canonical Wnt signaling (Figure 3E middle) and demonstrated decreased inflammatory responses (Figure 3F right), consistent with the broad immunosuppressive role of GAL1 in TME ^12^.

Stratification by PAM50 molecular subtype ^35^ revealed no statistically significant association with DFS within individual subtypes, although HER2-enriched tumors showed a trend toward significance (Figure S3D). The absence of statistical significance within subtypes likely reflects limited sample size following stratification rather than the absence of biological effect. To assess the impact of the gene pair ratio in a larger dataset, we used the online tool KMplotter to analyze relapse-free survival based on *LGALS1/ST6GAL1* expression. Comparing patients in the lowest (Q1) and highest (Q4) quartiles, we found that a high *LGALS1/ST6GAL1* ratio was consistently associated with worse relapse-free survival (RFS) across most breast cancer subtypes (Figure S3E). Thus, a combined expression profile characterized by high *LGALS1* and low *ST6GAL1* delineates a mesenchymal, immune-attenuated transcriptional program associated with poorer clinical outcomes in breast cancer.

### GAL1 deficiency delays early tumor development and metastatic dissemination in a spontaneous breast cancer model

Given that GAL1 critically regulates normal mammary branching morphogenesis (Figure 1), and that developmental programs are frequently co-opted during tumorigenesis ^1,2^, we investigated whether this glycan-binding protein influences the transition from normal to neoplastic mammary epithelium and whether it governs early events in tumor dissemination. We used the *MMTV-PyMT* spontaneous breast cancer model crossed onto a *Lgals1^-/-^* background (Figure S4A) to investigate the functional relevance of GAL1 in tumor initiation, progression, and metastasis. In this model, the polyoma middle T antigen (PyMT) is expressed in mammary epithelium under the *MMTV* promoter, driving stepwise progression from premalignant lesions to invasive carcinoma, with a high incidence of lung metastasis, recapitulating key features of human breast cancer ^37^ (Figure S4B). Interestingly, GAL1 deficiency in *MMTV-PyMT* x *Lgals1^-/-^* mice (Figure S4A, S4B) resulted in reduced tumor burden, evidenced by fewer transformed ducts and fewer detectable lesions in the mammary gland (Figure 4A). Recapitulating the phenotype observed in the normal gland (Figure 1), *PyMT Lgals1 ^-/-^* mice displayed a reduced frequency of PR⁺ cells per duct at 12 weeks of age (Figures 4B and 4C; Figures S4C-S4E). IHC of non-transformed MG confirmed GAL1 expression in the basal cell compartment (Figure 4D), and its absence was associated with a reduced MaSC fraction (Figure 4E). Remarkably, GAL1 deficiency delayed tumor onset resulting in considerably prolonged tumor-free survival (Figure 4F). Moreover, we also observed a substantial reduction in CD24^low^CD44^high^ cancer stem cells (CSC) both inside tumors and in the lung (Figure 4G and 4H). Importantly, lung metastasis was markedly impaired in GAL1-deficient mice, with reductions in both EpCAM^+^ PyMT^+^ disseminated tumor cells (DTC) in the lung and macrometastatic lesions (Figure 4I and 4J; Figure S4F).

**Figure 4:**
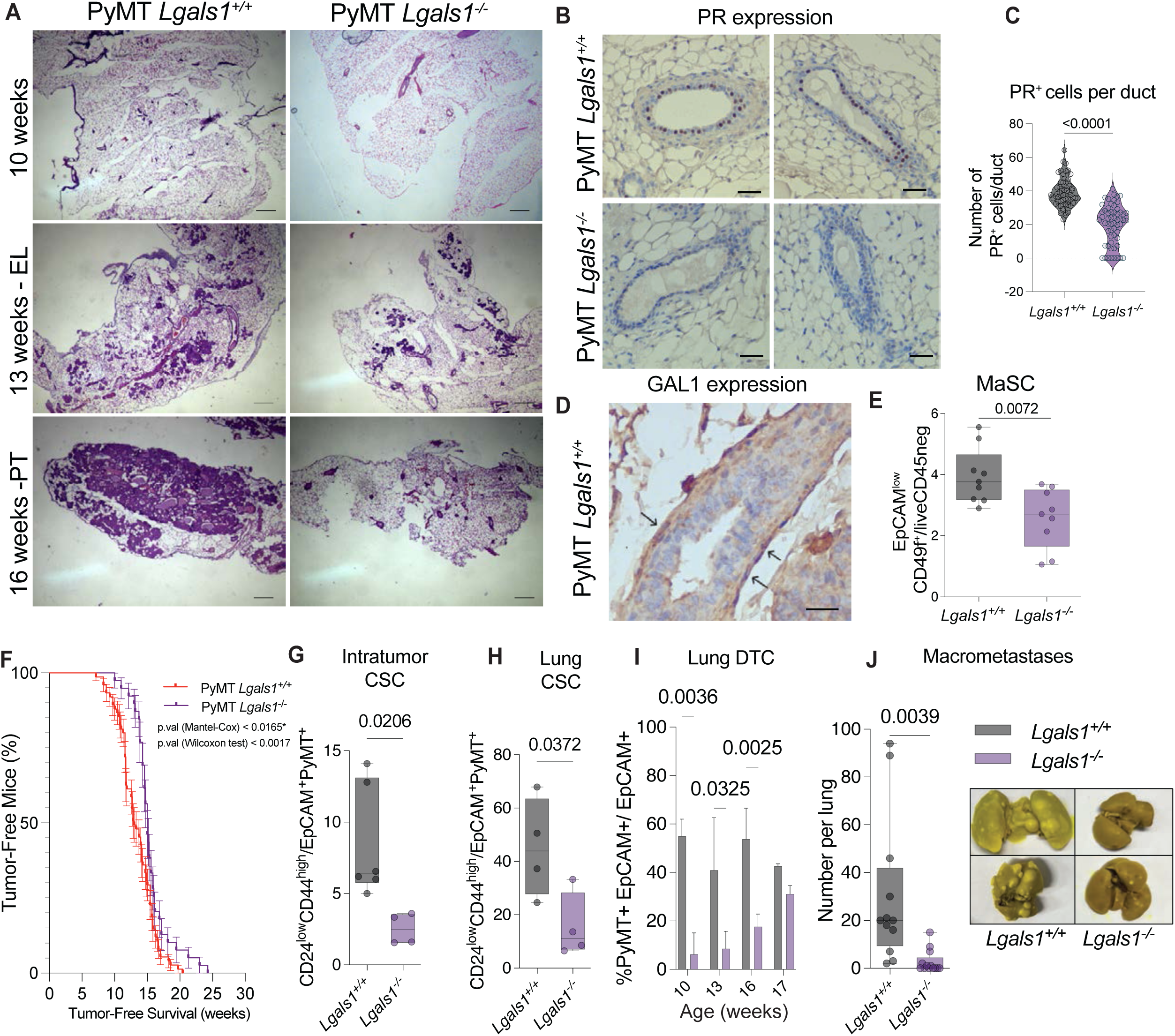
GAL1 deficiency impairs early dissemination in transgenic mice. A) Representative histological images of MG from *PyMT Lgals1^+/+^* and *Lgals1^-/-^*mice stained with hematoxylin and eosin (H&E) between weeks 10 to 16 (scale bar: 100 µm). B) Representative IHC images of PR^+^ nuclear staining (scale bar: 20 µm). C) Determination of PR^+^ cells per duct. Determination was performed by counting the number of PR^+^ nuclei relative to the total number of cells per duct. D) Detection and localization of GAL1 in mammary ducts of *PyMT* mice by IHC. Arrows indicate positive immunoreactivity in basal cells. E) Flow cytometry of MaSC in both *PyMT Lgals1^+/+^* and *Lgals1^-/-^* mice. F) Kaplan Meier curve showing tumor-free survival time in both *PyMT Lgals1^+/+^ and Lgals1^-/-^* mice. G-I) Flow cytometry of: G) mammary CSC in tumors of both *PyMT Lgals1^+/+^* and *Lgals1^-/-^*mice, H) CSC in lungs of both *PyMT Lgals1^+/+^* and *Lgals1^-/-^* mice, I) Epcam^+^PyMT^+^ DTC in lungs of both PyMT *Lgals1^+/+^*and *Lgals1^-/-^* mice at different ages. J) Determination of macrometastatic foci in lungs of both *PyMT Lgals1^+/+^* and *Lgals1^-/-^* mice at euthanasia, when all tumors had a similar volume. **Statistical analysis:** (C-E-G-H-I-J) Unpaired Welch’s t-test. p-values < 0.05 were considered statistically significant. All experiments were performed with n ≥ 4 mice per group (C) A total of 12 randomly selected ducts were analyzed per mouse sample. (F) Mantel-Cox and Gehan-BreslowWilcoxon test. n= 75 (*PyMT Lgals1^+/+^*), n= 25 (*PyMT Lgals1^-/-^*).

Our findings reveal that GAL1-dependent regulation of normal MG morphogenesis may be hijacked to drive early breast cancer onset and metastatic spread. The impaired expansion of MaSC and ER⁺ luminal progenitors, coupled with the reduced metastatic burden in *MMTV-PyMT Lgals1^-/-^* mice, highlights the therapeutic potential of targeting the GAL1-glycan axis to prevent tumor dissemination.

### Endogenous GAL1 accelerates early cancer cell dissemination in the spontaneous *MMTV-Neu* oncogene tumor model

Traditionally, metastasis has been viewed as a late event in cancer progression; however, accumulating evidence indicates that tumor cells can disseminate at very early stages of tumor development ^1,2^. These DTCs may exit the primary site while the tumor remains clinically undetectable and can either persist in a dormant state or contribute to metastatic outgrowth at later stages ^38^. Identifying the molecular cues that govern this early escape remains a major challenge. Building on these observations, we investigated whether GAL1 contributes to early cancer cell dissemination using the *MMTV-Neu* transgenic mouse model, which recapitulates key features of HER2⁺ human breast cancer and undergoes a slow progression from benign lesions to metastatic disease, thereby providing a temporal window to interrogate early tumorigenesis and dissemination before the emergence of bulky primary tumors (Figure 5A). We interrogated published transcriptomic profiles from early lesions (EL; 14 weeks), primary tumors (PT; 18 weeks), and early lesion-derived mammospheres, using RNA-seq datasets ^39^ (Figure 5B). HER2^+^ EL cells are engaged in a mesenchymal-like (M-Like) invasive program, whereas PT cells display a proliferative Epi-like phenotype ^39^. Notably, *Lgals1* emerged as the most upregulated galectin transcript in EL mammospheres relative to PT, while the sialyltransferase *ST6GAL1* was considerably downregulated (Figure 5B and 5C) at this stage.

**Figure 5:**
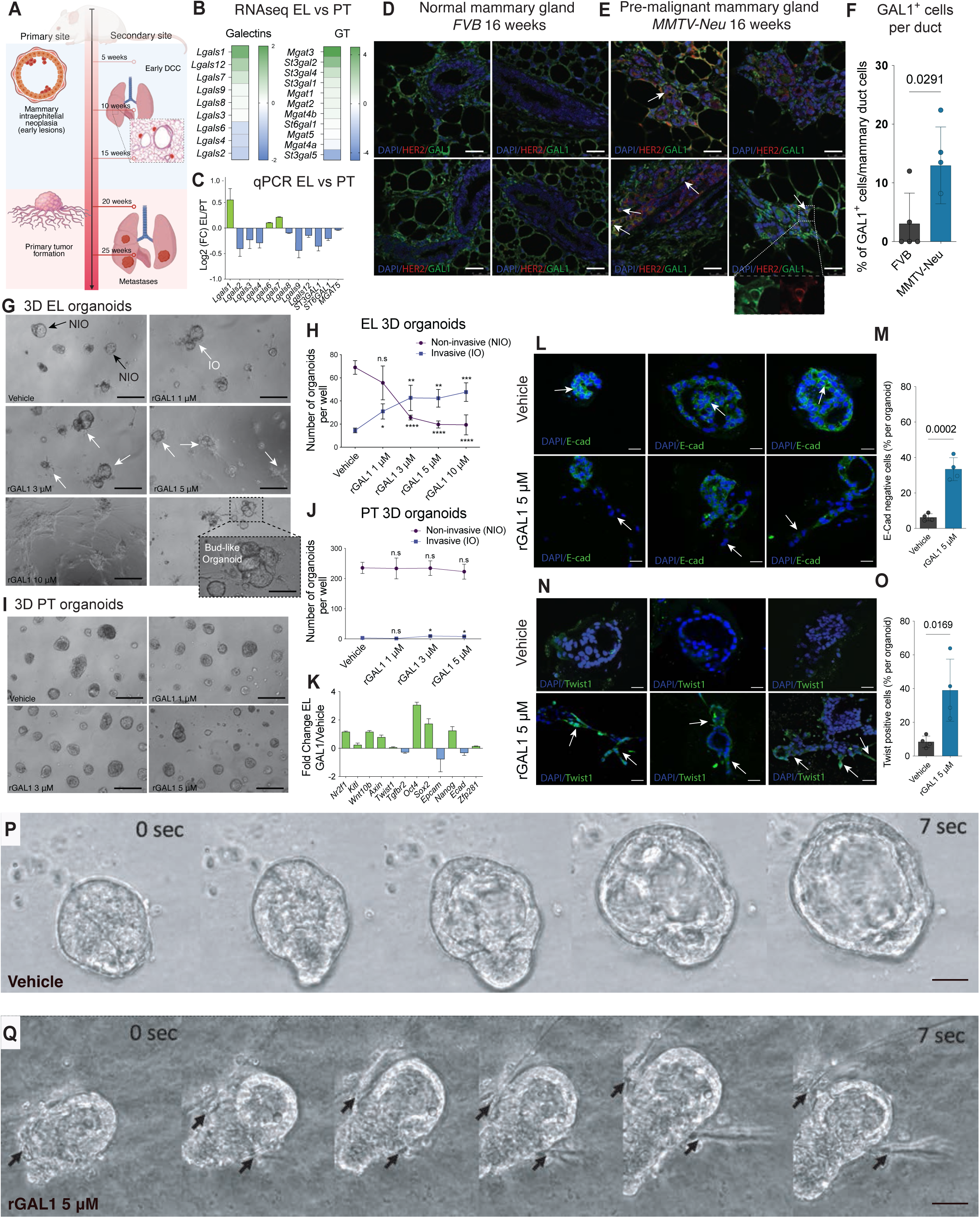
GAL1 triggers EMT and invasive phenotypes in EL MG 3D cultures. A) Schematic representation of different stages of the *MMTV-Neu* transgenic mouse model. B) Heatmap showing galectins and GT mRNA significantly expressed in MG 3D cultures, as determined by RNAseq (EL/PT ratio) (FDR < 0.05 and FC > 2 or < 0.5).. C) RT-qPCR validation of galectins and GT expression in MG 3D cultures (EL/PT ratio). D-E) Immunofluorescence images showing GAL1 staining in normal FVB MG tissue (scale bar: 25 µm) (D) and in *MMTV-Neu* EL (14 weeks) (E). F) Determination of GAL1^+^ cells per duct in both normal and pre-neoplastic MG. Each dot represents the average of GAL1^+^ cells per duct per MG per mice. G) Microscopy images of EL 3D cultures treated for 7 days with different concentrations of recombinant GAL1 (rGAL1) or vehicle control (scale bar: 50 µm). H) Determination of invasive and non-invasive organoids following rGAL1 treatment of EL derived 3D cultures. I) Microscopy images of PT 3D cultures treated with different concentrations of rGAL1 for 7 days or with vehicle (scale bar: 50 µm). J) Determination of invasive and non-invasive organoids of PT. K) Heatmap of genes related to EMT and the Wnt pathway determined by RT-qPCR (rGAL1 treated/untreated ratio) in EL. L) Immunofluorescence of the epithelial cell marker E-cad in 3D cultures treated with vehicle or rGAL1 M) Determination of E-cad-negative cells. Each dot represents the average of E-cad-negative cells per organoid per replicate. N) Immunofluorescence of the mesenchymal marker Twist1 in 3D cultures treated with rGAL1 or vehicle. O) Determination of Twist1^+^ cells is shown. Each dot represents the average of Twist1^+^ cells per organoid per replicate. P-Q) Time-lapse imaging (24 h) of EL organoids treated with vehicle (P) or rGAL1 (Q) (Scale bar: 10 µm) **Statistical analysis**: (F-H-J-M-O) Unpaired Welch’s t-test (J). p-values < 0.05 are considered significant. All experiments were performed with n = 4 mice per group.

We validated these observations in age-matched controls and *MMTV*-Neu mammary glands at the EL stage (14 weeks). In normal glands, GAL1 showed periductal and stromal expression. In pre-neoplastic lesions (EL) we observed an increased frequency of GAL1^+^ cells including cells coexpressing GAL1 and HER2 (Figure 5D, 5E and 5F). To functionally assess the effect of exogenous GAL1, we treated 3D Matrigel organoids derived from EL lesions with recombinant GAL1 (rGAL1). rGAL1 induced a dose-dependent invasive phenotype (Figure 5G and 5H), with cells escaping the organoid body and extending mesenchymal-like protrusions and bud-shaped structures reminiscent of TEB-like structures observed during branching MG morphogenesis. In contrast, PT organoids were refractory to GAL1-induced invasion, indicating a stage-restricted sensitivity to this lectin (Figures. 5I and 5J). To gain mechanistic insights into this phenotypic transition, we analyzed expression of a set of genes associated with this mesenchymal phenotype in organoids from rGAL1-treated EL. We observed an increase in the expression of EMT- and stemness-associated genes such as *Oct4, Sox2* and *Nanog* as well as Wnt/β-catenin associated genes such as *Axin*, *Wnt10b* and *Twist1*, and a decrease in the epithelial cell markers *EpCAM* and *E-Cadherin (E-Cad)* (Figure 5K). These transcriptional changes were corroborated by immunofluorescence analyses of 3D cultures. Indeed, rGAL1 reduced the percentage of E-Cad^+^ cells (Figure 5L and 5M) and increased the frequency of Twist1^+^ cells (Figure 5N and 5O). Time-lapse microscopy visualized dynamic cell egress from the 3D organoids following GAL1 exposure (Figure 5P and 5Q). Thus, GAL1 functions as a pro-invasive cue during normal development, which transformed cells may co-opt at early neoplastic stages to promote dissemination.

### GAL1 promotes EMT and stem-like phenotypes in human HER2^+^ breast cancer cells

Given the up-regulation of GAL1 in early HER2⁺ lesions in the *MMTV*-*Neu* model and its ability to induce EMT/stemness-associated transcriptional and phenotypic changes in organoids derived from early lesions, we next sought to explore whether these mechanisms are conserved in human HER2⁺ breast cancer cells.

We selected two human HER2⁺ breast cancer cell lines with distinct aggressiveness profiles ^40^ (Figure S5A). BT-474 is a HER2^+^ breast poorly aggressiv cancer cell line responsive to trastuzumab (Tz), while JIMT-1 exhibits basal-like properties, is highly aggressive and resistant to Tz ^41^. JIMT-1 showed higher GAL1 mRNA and protein expression (Figure 6A, 6B and 6C) than BT-474, greater cell surface GAL1 binding (Figure 6D), and lower *ST6GAL1* expression and SNA binding (Figure 6E and 6F), consistent with superior GAL1 activity associated with this weakly sialylated phenotype (Figure 6G).

**Figure 6:**
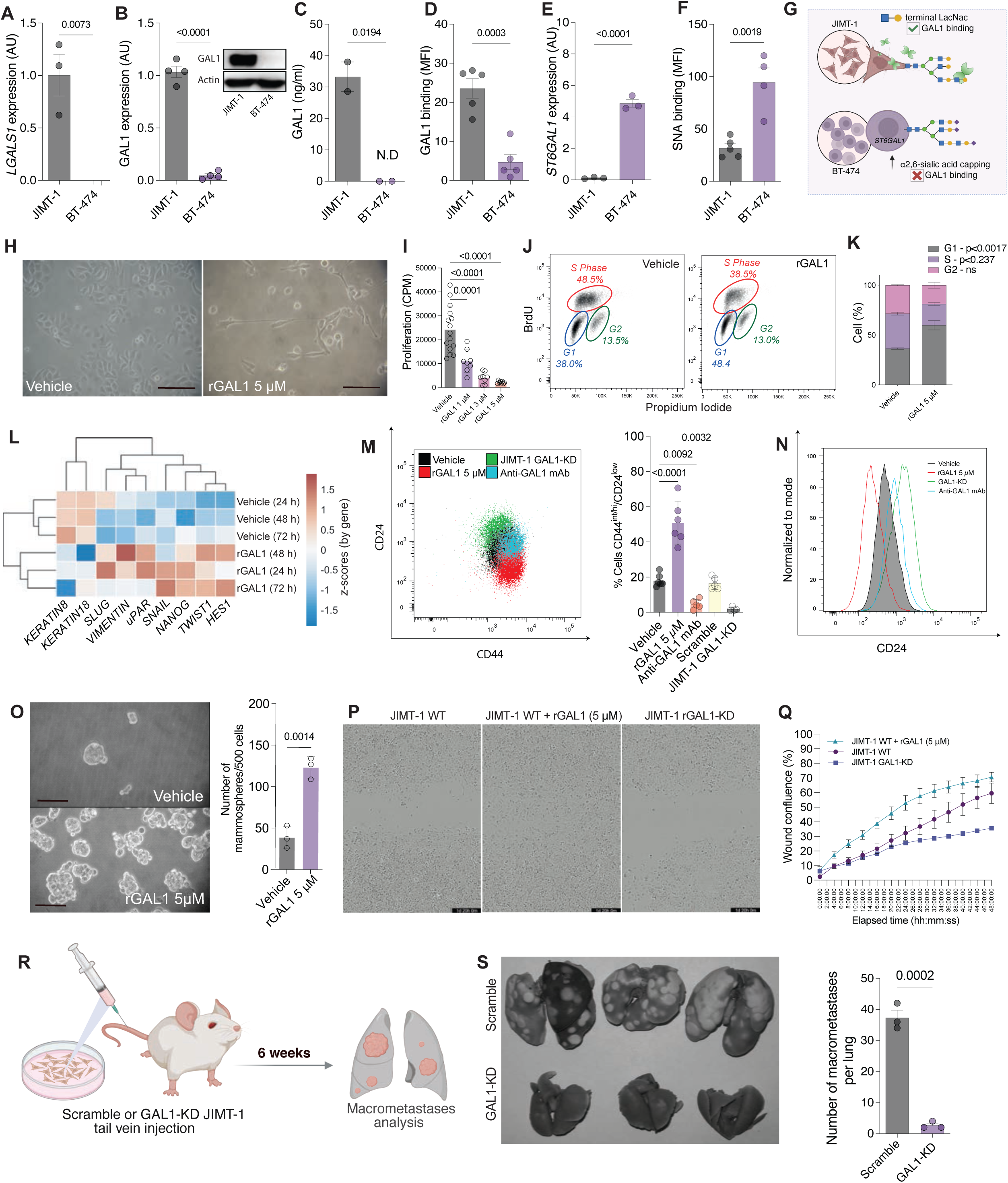
GAL1 promotes EMT and cancer stem cell phenotypes in human breast cancer cells. A-B) *LGALS1* mRNA (A) and GAL1 protein (B) expression in JIMT-1 and BT-474 cell lines. C) GAL1 protein concentrations in JIMT-1 and BT-474 cell lines CM (ELISA). D) GAL1 binding to JIMT-1 and BT-474 cell lines (flow cytometry). E) *ST6GAL1* mRNA expression in JIMT-1 and BT-474 cell lines (qPCR). F) SNA binding to JIMT-1 and BT-474 cell lines (flow cytometry). G) Schematic representation of analyzed cell lines summarizing their glycophenotype. H) Bright-field images of JIMT-1 cells treated with vehicle or rGAL1 (5 µM, 72 h) (Scale bar: 50 µm). I) Proliferation, measured by ^3^ [H]-thymidine incorporation after addition of rGAL1 (1, 3 or 5 µM) in JIMT-1 cells. J) Cell cycle analysis (propidium iodide plus +BrdU) by flow cytometry). K) Determination of the G1/ S /G2 phases. L) Heatmap of EMT and stemness markers (RT-qPCR, normalized). M) Dot plot of JIMT-1 cells treated with rGAL1 (5 µM) stained for CD24 and CD44 and analyzed by flow cytometry (left panel), percentage of CD24^low^/CD44^high^ cells across different experimental conditions (right panel). N) Flow cytometry of CD24 normalized to mode (Scale bar: 100 µm). O) Mammosphere formation: cells pre-treated with rGal-1 (5 µM, 72 h), spheres counted per 500 seeded cells. P) Wound-healing assay Q) Wound confluency over 48 h. R) Schematic representation of the tail vein injection experimental design. S) Lung images and assessment of macrometastases at 6 weeks in KD or scramble JIMT-1-injected-mice. **Statistical analysis:** (A-B-C-D-E-F-O-S) Unpaired Welch’s t-test. P-values < 0.05 were considered significant. (I-M) One-way ANOVA with Dunnet’s post-test (vs Vehicle). All experiments were performed with n ≥ 3 per group.

Treatment of JIMT-1 cells with rGAL1 led to a striking morphological switch toward a mesenchymal-like phenotype, similar to that observed in organoids from the *MMTV-Neu* model (Figure 6H). JIMT-1 cells also exhibited a dose-dependent inhibition of proliferation (Figure 6I; Figure S5B), accompanied by a cell cycle arrest in G1 without any sign of early apoptosis (Figure 6J and 6K; Figure S5C). The less aggressive cell line BT-474, displaying a GAL1-restrictive glycophenotype, did not undergo this quiescent-like shift (Figure S5D).

Furthermore, GAL1 increased the expression of EMT-related (*TWIST, HES, SNAIL, SLUG, VIMENTIN*) and stemness-related (*NANOG* and *uPAR*) genes (Figure 6L; Figure S5E). Additionally, rGAL1 treatment increased the frequency of CD24^low^/CD44^+^ CSC-enriched population in JIMT-1 cells (Figure 6M and 6N; Figure S5F, S5G). As observed in 3D-organoids (Figure 5G), we also found a marked reduction of E-cad, indicating a shift towards a mesenchymal identity (Figure S5H). Knocking down GAL1 using shRNA strategies in the JIMT-1 cell line (GAL1-KD) or blocking GAL1 with a specific neutralizing monoclonal antibody (mAb) ^14^ (Figure 6M, 6N and Figure S5I), reduced the CD24^low^/CD44^+^ CSC-enriched phenotype, reinforcing the specificity and therapeutic relevance of targeting the GAL1-glycan axis. Consequently, rGAL1 increased mammosphere formation (Figure 6O) and enhanced migration in scratch-wound (Figure 6P and 6Q) and transwell migration (Figure S5J and S5K), further supporting its role in driving a migratory and invasive phenotype.

To further validate these findings *in vivo*, we performed tail vein injections of JIMT-1 GAL1 WT or GAL1-KD cells into immunocompromised *Scid/NOD/IL2R^-/-^* mice and monitored metastatic colonization (Figure 6R). GAL1-silenced JIMT-1 cells failed to generate macrometastasis, underscoring the contribution of this glycan-binding protein to a successful metastatic colonization (Figure 6S). Together, these results show that GAL1 is associated with EMT-related changes, stem-like features, and metastatic behavior in aggressive HER2⁺ human breast cancer cells.

### Antibody-mediated GAL1 blockade impairs MG branching morphogenesis and reduces metastatic dissemination

We next studied whether systemic GAL1 neutralization may restrain early escape and metastasis. For this, *MMTV-Neu* mice were treated with the anti-GAL1 neutralizing (mAb) ^13,14^ beginning at 14 weeks of age (EL stage) and continuing through week 19. Mice were euthanized at week 25 to assess primary and metastatic outcomes (Figure 7A).

**Figure 7:**
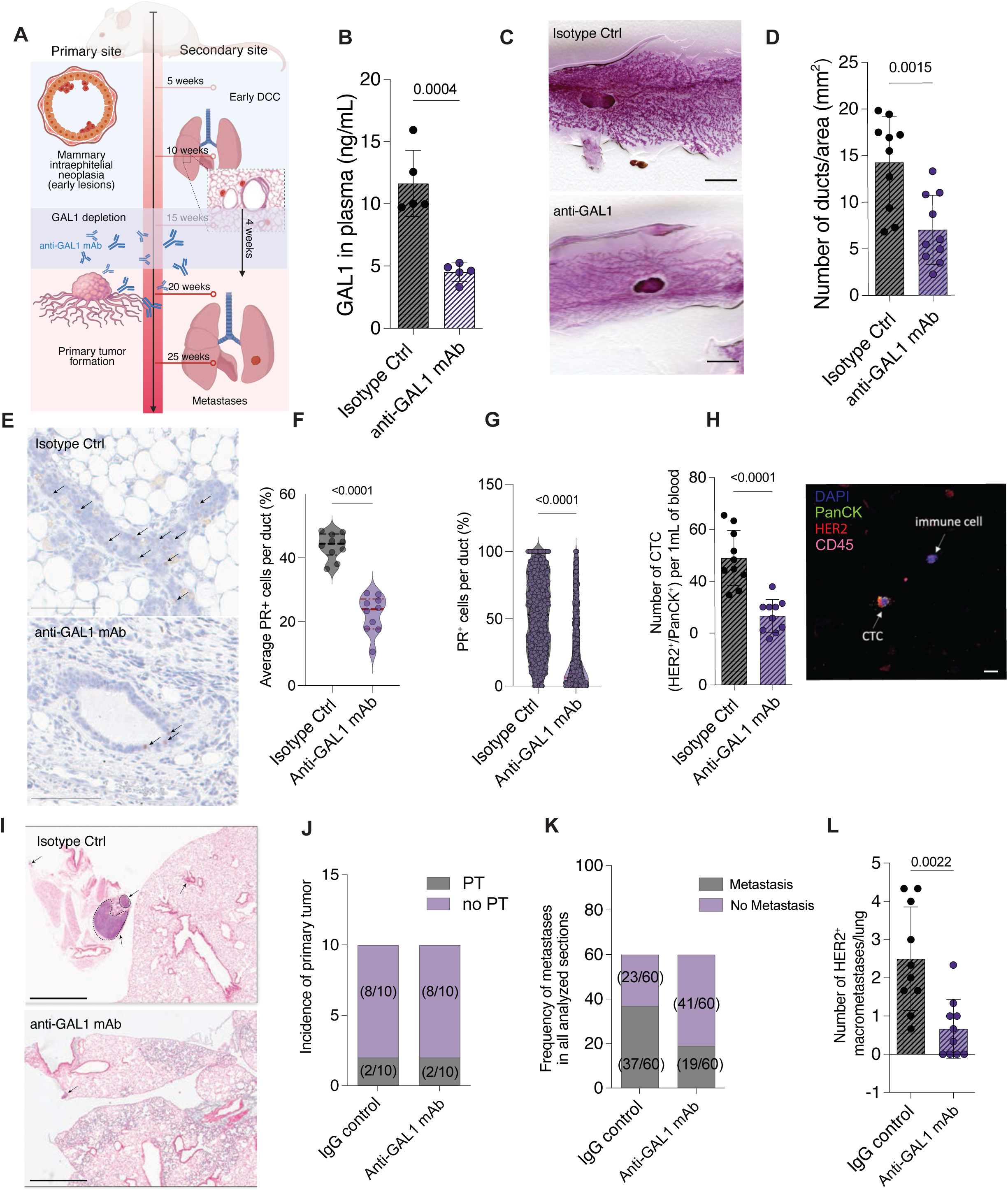
**Systemic GAL1 neutralization reduces early dissemination in *MMTV-Neu* mice**. A) Scheme of the therapeutic strategy used in the *MMTV-Neu* mouse model. Mice were treated with anti-GAL1 mAb or isotype control (30 mg/kg/mouse) twice a week (for 4 weeks). B) Circulating GAL1 in blood (ELISA). C) Representative mammary whole mounts of treated mice (Scale bar: 5 mm). D) Determination of the number of ducts per area of MG in response to anti-GAL1 mAb or isotype control treatment. E) Representative microscopy images of PR staining in *MMTV-Neu* MG (Scale bar: 100 µm). F) Determination of the number of PR^+^ cells per MG. G) Determination of the number of PR^+^ cells per duct. H) Determination of the number of CTC in blood before euthanasia (Scale bar: 20 μm). I) Representative microscopy images of lungs sections stained with H&E. Arrows indicate macrometastatic foci (Scale bar: 1 mm). J) Incidence of PT tumors per group. K) Frequency of macrometastases per lung section. L) Determination of the number of macrometastases per lung. **Statistical analysis:** (B-D-F-G-H) Unpaired Welch’s t-test (J). Unpaired Mann-Whitney’s t-test. P-values < 0.05 are considered significant. All experiments were performed with n > 4 mice per group.

Treatment with the anti-GAL1 mAb markedly reduced circulating GAL1 levels (Figure 7B), indicating effective systemic target engagement. Whole-mount analysis revealed a decreased branching morphogenesis in the MG of anti-GAL1-treated mice, mirroring the phenotype observed in *Lgals1^-/-^* mice (Figure 7C). Histopathological analysis further demonstrated reduction in the numbers of transformed ducts (Figure 7D). Given the ability of breast cancer cells to hijack progesterone-driven developmental programs that control branching to disseminate^2^ and considering the loss of PR^+^ cells induced by GAL1 in normal and *PyMT* glands (Figures 1 and 4), we examined whether therapeutic GAL1 neutralization alters PR abundance. As observed in both C57BL/6 *Lgals1^-/-^*and *PyMT Lgals1^-/-^* mice, depletion of GAL1 altered PR expression in the MG, dampening the number of PR^+^ nuclei per duct and per MG (Figure 7E, 7F and 7G), further suggesting the ability of GAL1 to regulate the progesterone-PR axis during normal breast morphogenesis and in early breast cancer progression.

Importantly, GAL1 neutralization markedly reduced the number of CTC (HER2⁺/PanCK⁺) in peripheral blood (Figure 7H), suggesting that early dissemination was impaired in response to therapeutic GAL1 depletion. This reduction was accompanied by a decrease in lung metastatic burden (Figure 7I). Whereas primary tumor incidence was comparable between groups (Figure 7J), both macrometastases frequency (Figure 7K) and number significantly decreased upon anti-GAL1 mAb treatment (Figure 7L).

In summary, early systemic GAL1 neutralization reduced MG branching morphogenesis, mitigated CTC release, and diminished lung metastatic burden, providing preclinical evidence that GAL1 contributes to early dissemination and metastatic outgrowth, and supporting GAL1-glycan interactions as a tractable target to limit metastatic progression in breast cancer.

## DISCUSSION

Metastatic breast cancer remains the leading cause of cancer-related death in women worldwide (https://seer.cancer.gov), underscoring the need for new therapeutic strategies that directly target metastatic progression. Increasing evidence indicates that dissemination occurs early during breast tumorigenesis ^1,2^, highlighting the importance to identify molecular pathways that enable these early dissemination stages. Here, we uncover a shared mechanistically linked role of GAL1-glycan interactions in orchestrating MG development and facilitating early metastatic dissemination, revealing a novel therapeutic vulnerability. Specifically, we identify GAL1, an evolutionarily conserved glycan-binding protein, as a key player in MG morphogenesis that is co-opted during tumorigenesis. In the normal gland, GAL1 is enriched in basal and MaSC and supports progesterone-driven branching morphogenesis. Loss of GAL1 impairs ductal growth and reduces the MaSC pool. Moreover, GAL1 was associated to a lower frequency of ER⁺ luminal cells and attenuated progestin responses.

Although the causality of this physiopathological transition, either direct or indirect, mediated by changes in epithelial hierarchy, remains to be further scrutinized, these findings are consistent with a role of GAL1 as a mediator of the hormonal response in MG. Interestingly, previous work has shown that nuclear translocation of GAL1, which promotes branching and invasion, is regulated by glycosylation. Specifically, α2,6-sialylation of LacNAc motifs releases GAL1 for nuclear import, whereas unsialylated glycans retain this lectin at the extracellular compartment ^24^. Our findings extend this framework, by showing that glycosylation-dependent gating operates alongside hormonal signals, reinforcing the concept of the glycan code governing GAL1 function. The marked reduction in PR expression observed in *Lgals1*-deficient mice across multiple complementary approaches suggests that GAL1 is required to maintain appropriate PR expression *in vivo*. Given the central role of PR signaling in progesterone-driven mammary morphogenesis and mammary stem cell development, diminished PR expression likely contributes to the defective nulliparous and MPA-induced branching and MaSC expansion observed in the absence of GAL1. Mechanistically, our data provide evidence that GAL1 localizes not only to the cytoplasm but also to the nucleus and chromatin, where it associates with *PGR-A* and *-B* promoters. Although additional studies will be required to define the molecular partners through which GAL1 may influence *PGR* transcriptional regulation, these findings support a model in which chromatin-associated GAL1 may directly or indirectly influence PR expression. These findings further support an emerging role of GAL1 in transcriptional regulation beyond its well-established extracellular glycan-dependent and intracellular signaling roles, suggesting that regulation of hormone receptor expression may represent an additional mechanism by which GAL1 influences MG development and stem cell function. In this regard, we have recently identified a nuclear role for GAL1 in regulating the transcription of oncogenes such as KRAS in pancreatic stellate cells (PSC), a key component of the aggressive pancreatic ductal adenocarcinoma (PDAC) ^42^. In those cells, GAL1 plays a critical role driving PSC activation and promoting the secretion of pro-tumorigenic cytokines and growth factors^42^.

The relative contribution of extracellular versus intracellular GAL1, including its nuclear shuttling remains to be defined in native tissue. Interestingly, flow-cytometric analysis of luminal compartments revealed an imbalance in ER⁺ luminal progenitors in GAL1-deficient MG. Because ER^+^ luminal progenitors are highly proliferative and elongate and migrate within TEB ^43^; the reduction of this compartment in *Lgals1*^-/-^ glands may likely contribute to diminished branching and may cooperate at a further stage to delay tumor onset ^44^, emphasizing a molecular interconnection between MG branching and early tumor establishment.

Consistent with a role of breast cancer cells in co-opting the GAL1-glycan pathway for early tumorigenesis and dissemination, genetic ablation or therapeutic blockade of this lectin in two spontaneous models (*MMTV-PyMT* and *MMTV-Neu)*, reduced transformed ducts, and decreased lung metastases and CTC, highlighting the relevance of GAL1 as a multifunctional therapeutic target. Functionally, GAL1 promoted EMT and stemness signatures (*Twist, Snail, Vimentin; Nanog, Oct4, Sox2*) in mouse organoids and human HER2⁺ cells. In the *LGALS1^high^/ST6GAL1^low^* JIMT-1 cell line, rGAL1 induced a quiescent, migratory, EMT-like state (CD24^low^/CD44^high^) and enhanced sphere formation and motility, whereas GAL1 silencing curtailed metastatic colonization *in vivo*. These observations are consistent with reports implicating GAL1 in EMT through TGF-β/Smad and Hedgehog pathways in other epithelial cancers, suggesting a conserved role for this lectin in promoting EMT ^17,45^.

Because terminal α2,6-sialylation interrupts GAL1-glycan binding, we examined whether the *LGALS1*/*ST6GAL1* balance has clinical prognostic relevance. Across breast cancer datasets, a *LGALS1*^high^*/ST6GAL1^low^*expression signature was associated with poorer clinical outcomes in patients and was characterized by enrichment in EMT, canonical *Wnt* and immune evasion signatures. These findings raise the possibility that tumors with a *LGALS1*^high^/*ST6GAL1*^low^ profile may be particularly dependent on GAL1 signaling and could therefore be targeted with anti-GAL1 mAb. Consistently, it has been shown that chemotherapy-induced *ST6GAL1* loss in TNBC promotes CTC clustering and metastatic seeding ^22^. This suggests that loss of *ST6GAL1* may create a permissive glycophenotype that enhances GAL1-stroma interactions, further driving metastasis. While GAL1’s prognostic value is well established in colon ^13,15^, PDAC ^16,46^ and myelofibrosis ^47^, the role of *ST6GAL1* is more complex, with context-dependent associations requiring tumor-specific analysis of this axis.

Importantly, systemic administration of an anti-GAL1 mAb in the *MMTV-Neu* model recapitulated GAL1-deficient phenotypes, reducing PR⁺ cells, early lesions, DTC, and lung metastasis. The reversion of the *MMTV-Neu* aggressive phenotype when GAL1 was neutralized was associated to a drop in CTC and DTC counts. This *in vivo* effect is supported by evidence showing that progesterone-driven signals promote early dissemination of HER2^+^ cells ^1,2^. Thus, lowering PR⁺ cells and subsequent progesterone signaling via GAL1 neutralization, may blunt early escape in breast cancer. These findings suggest that, at least in our models, GAL1 plays a key role at the initial steps of dissemination and escape, rather than influencing primary tumor growth, since reduction in CTC and metastasis occur in the absence of marked changes in primary tumor incidence. Because fibroblasts and immune cells also express GAL1 ^13^, an important component of the antimetastatic benefit may arise from effects on stromal and immune compartments. Beyond direct GAL1 inhibition, strategies to modulate the associated glycosylation machinery also hold promise, with the potential to block dissemination and enhance therapeutic responses.

Finally, while GAL1’s role as an immune regulator is broadly documented ^13,15,16,19,47,48^, our work reveals its contribution to both mammary branching and early dissemination, two mechanistically related processes. Given that both intracellular and extracellular GAL1 pools may likely contribute to tumor dissemination, dissection of the cellular source of this lectin and its subcellular compartmentalization add additional layers of complexity to its function. Although basal and myoepithelial lineages appear to be the predominant sources of GAL1 in normal MG, fibroblasts, tumor-associated immune cells, and cancer cells themselves may secrete GAL1 during malignant progression, creating overlapping; yet distinct glycan-dependent signaling niches. Tumor glycosylation patterns, particularly α2,6-sialylation controlled by ST6GAL1, further shape these extracellular effects. This represents an important axis of TME communication, as ST6GAL1 not only modulates GAL1-glycan interactions but also influences the behavior of key stromal and immune cell populations through diverse mechanisms including the creation of Siglec-specific glyco-epitopes ^49^. Interestingly, previous studies demonstrated a key role for ST6GAL1 in acinar to ductal metaplasia, a process that fosters tumorigenesis in PDAC ^50^. Moreover, we demonstrated that ST6GAL1-dependent sialylation controls GAL1-driven induction of myeloid-derived suppressor cells ^13^, establishment of angiogenesis programs ^18,51^, and T helper cell polarization ^52^. Future studies are warranted to investigate how GAL1-glycan interactions modulate the interplay between PR⁺ luminal cells and immune cells populations within the pre-neoplastic gland and metastatic niches, and how distinct epithelial, stromal, and immune compartments contribute to GAL1 secretion and function. Together, these insights will provide a framework to understand how glycosylation-driven signaling networks orchestrate early tumor evolution and shape the immunosuppressive landscape of breast cancer.

Thus, GAL1 emerges as a key regulator of hormone-driven mammary development and a critical determinant of early metastatic dissemination. By mechanistically linking GAL1 to PR activity, cancer cell stemness and EMT, and integrating these processes within an ST6GAL1-controlled glycosylation program, our findings establish the GAL1-glycan axis as a biologically grounded and therapeutically tractable vulnerability in breast cancer. Given its interconnected roles in developmental plasticity, early dissemination, and immune regulation, targeting GAL1 or therapeutically rewiring receptor glycosylation may complement existing treatments and improve outcomes for patients with metastatic breast cancer.

## ACKNOWLEDGEMENTS

We thank all members of M.S, G.A.R and J.A.G labs for intellectual insights and support. We thank Mirta Schattner for critical reading of the manuscript and Claudia Lanari (IBYME) for the *NOD/LtSz-scid/IL-2Rgammanull* mice. This work was supported by grants from Agencia de Investigación, Desarrollo e Innovación (PICT 2014-3687 and 2017-0494 to G.A.R and PICT 2018-02602 and 2020-00874 to M.S.), Ministerio de Ciencia, Tecnología e Innovación (Redes Federales de Alto Impacto) to G.A.R., Bunge & Born, Sales, Baron and Richard Lounsbery Foundations to G.A.R and a visiting fellowship in J.A.G lab from Fundacion Sales to R.M.P. We thank the Ferioli, Ostry, Caraballo and Alfonzo families for kind donations. P.N is funded by MICIU/ISCIII-ERDF grant PI23/00591. L.E.V.S was funded by a Medical Scientist Training Program grant from the National Institute of General Medical Sciences (NIGMS) of the NIH under award number T32GM149364 to the Albert Einstein College of Medicine MD-PhD Program. J.A.G. is supported by grants from The National Institute of Health (NIH) /National Cancer Institute (NCI) (CA109182-23, CA284085-1 and CCSG P30 CA013330-52), and The Gurwin Foundation, DOD CDMRP FY23 BCRP FL2 PSPS (BC230537-HT94252410078) and an Aging and Cancer grant from the Samuel Waxman Cancer Research Foundation and The Mark Foundation for Cancer Research. Schematic representations were created with BioRender.com. We also acknowledge the Core Facilities at MECCC and AECOM including Flow Cytometry, Institute for Animal Studies, Analytical Imaging Facility and the Gruss Lipper Biophotonics Center. The cores at MECCC are funded by the NCI Cancer Center Support Grant P30 CA CA013330-52. We also acknowledge the use of the animal research facilities at the Albert Einstein College of Medicine (AECOM) and the Analytical Imaging Facility at AECM. AECM facilities are funded through NIH SIG #1S10OD026852-01A1.

## AUTHOR CONTRIBUTIONS

Conceptualization, R.M.P., G.A.R. and M.S.; Methodology and Investigation, R.M.P., M.B., L.E.V.S., T.D.M., Y.D.M., B.V.P, N.M.B, J.A.A.G., G.A.R. and M.S.; Resources, R.M.P., R.M.M., S.G.G., J.M.P.S., J.A.A.G.; Formal Analysis, R.M.P., M.B., L.E.V.S., T.D.M., Y.D.M., P.N., R.B., J.A.A.G., G.A.R. and M.S.; Data Curation R.M.P., M.B., L.E.V.S., Y.D.M.; Supervision, Project Administration and Funding Acquisition, G.A.R. and M.S.; Visualization, R.M.P., J.A.A.G., G.A.R. and M.S.; Writing Original Draft, R.M.P., G.A.R. and M.S.; Writing Review & Editing, all authors.

## RESOURCE AVAILABILITY

### Lead contact

Further information and requests for resources and reagents should be directed to and will be fulfilled by the Lead Contact: Mariana Salatino or Ramiro M. Perrotta.

### Materials availability

This study did not generate new unique reagents.

### Data and code availability

This paper analyzes existing, publicly available data. Accession numbers for the datasets are listed in the Key Resources Table. This paper does not report new original codes.

---

**Figure S1:**
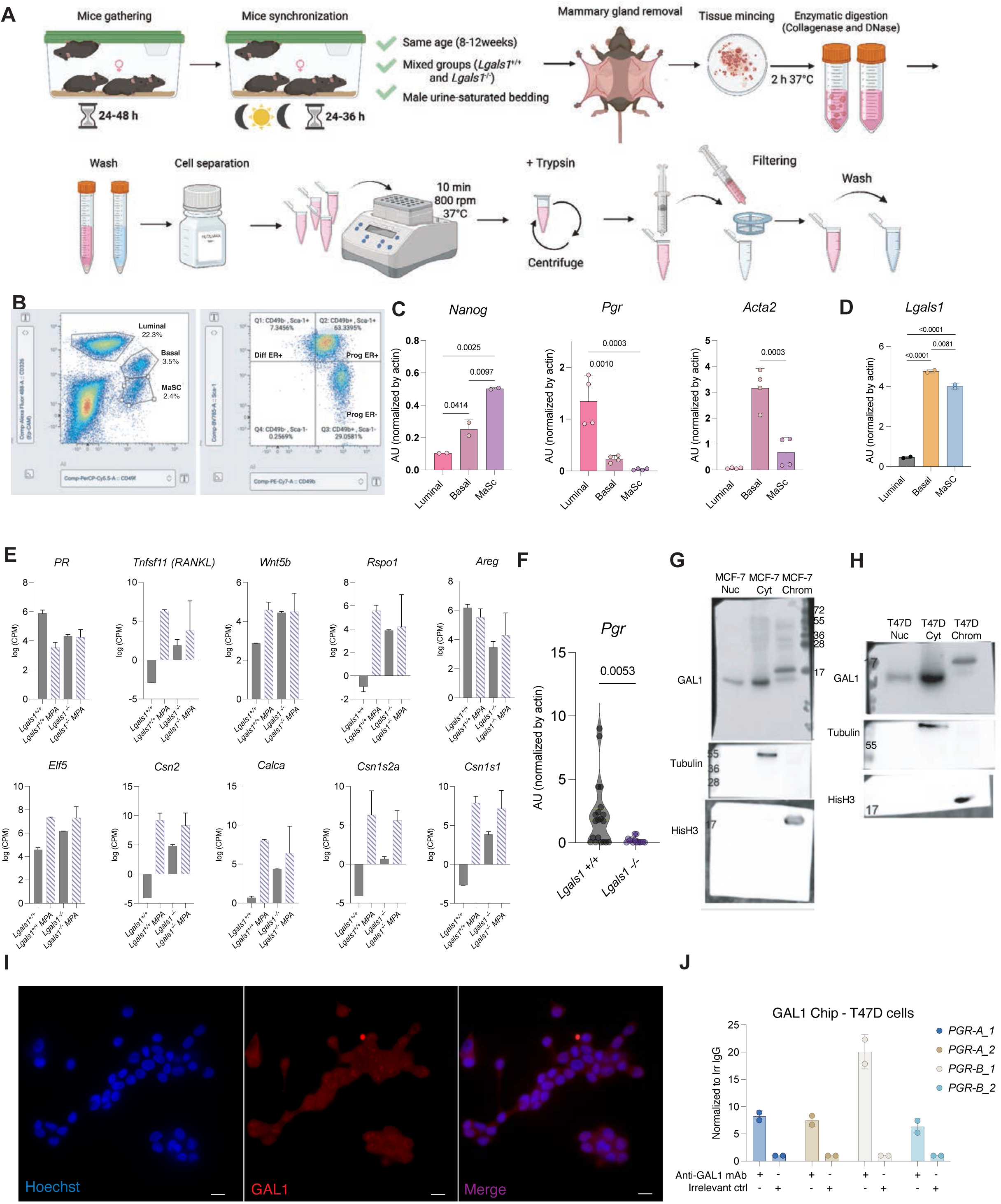
**Characterization of MG epithelial lineage**. A) Schematic workflow for isolation and characterization of the epithelial cell lineage in C57BL/6J MG. B) Representative dot plot showing the flow cytometry gating strategy. Q1: Diff ER^+^, Q2: Prog ER^+^ Q3: Prog ER^-^. C) mRNA analysis of lineage-specific markers to confirm the identity of each cell type (qPCR). D) GAL1 expression across the different MG epithelial cell lineages (qPCR). E) Expression of MG developmental and functional markers (RNAseq). F) PR mRNA quantification across *Lgals1^+/+^* and *Lgals1^-/-^* MG. G-H) Western blot analysis of GAL1 in cytoplasmic and nuclear fractions of MCF-7 (G) and T47D (H) cells. Fraction purity was assessed using compartment-specific markers (tubulin for the cytoplasm and histone-H3 for the chromatin fraction). Representative blots from n = 3 independent experiments are shown. I) Representative immunofluorescence images showing GAL1 expression (red) in MCF-7 breast cancer cells. Nuclei were counterstained with DAPI (blue). Scale bar, 20 μm. J) ChIP-qPCR validation of GAL1 enrichment at progesterone receptor (*PGR*) genomic regions. Four primer pairs spanning *PGR* regulatory regions (two targeting *PGR-A* and two targeting *PGR-B*) were used to assess enrichment. **Statistical analysis:** (J) Data are presented as mean ± SD from 2 independent experiments. (F) Unpaired Welch’s t-test. p-values < 0.05 are considered significant.

**Figure S2:**
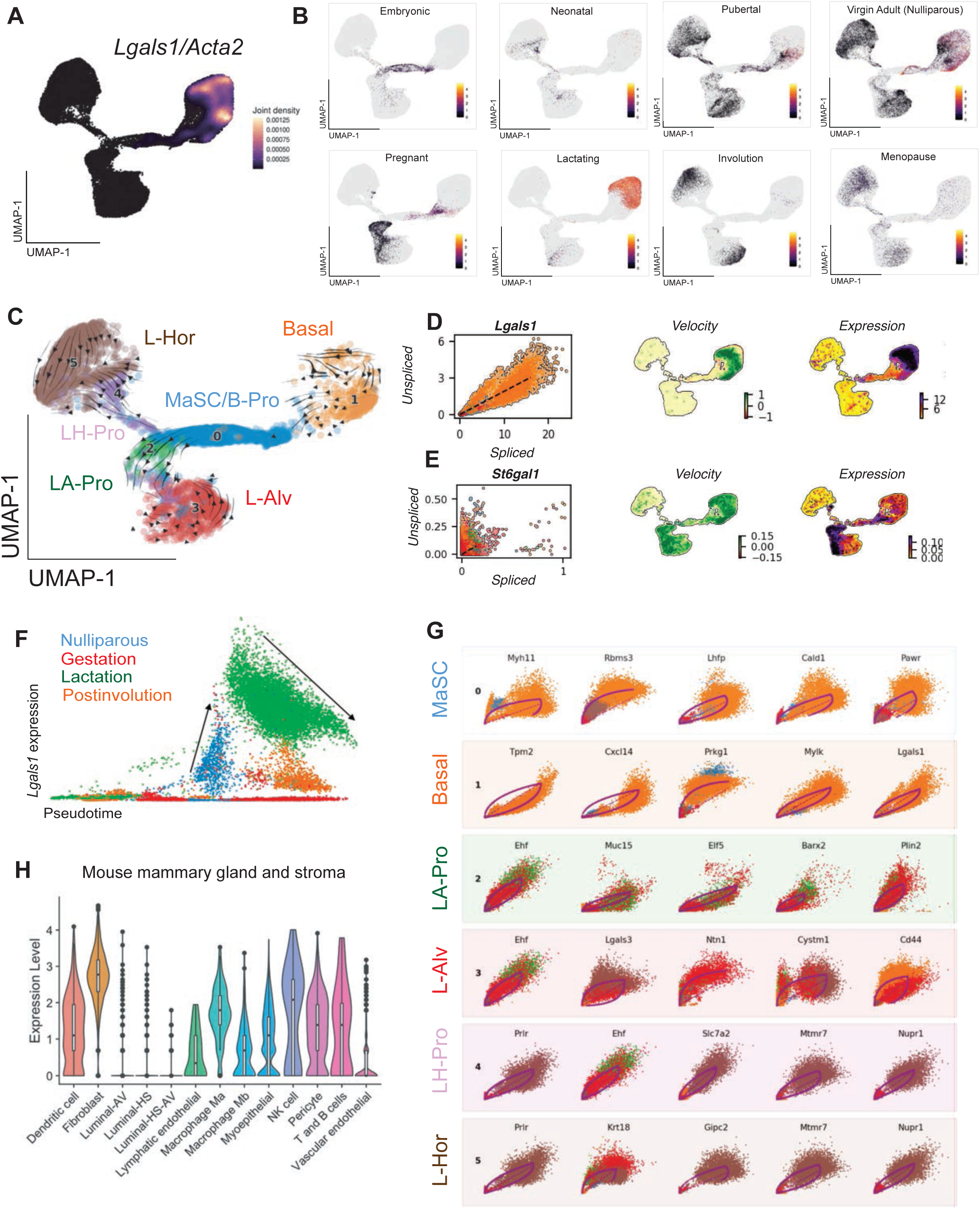
scRNAseq and velocity analysis of mouse MG. A) Visualization of *Lgals1* and *Acta2* (joint density) expression visualized on the 2D UMAP across the mouse MG ATLAS. B) *Lgals1* expression visualized on the 2D UMAP across distinct mammary cell lineages at each developmental stage of the MG. C) The velocities derived from scRNAseq data from mouse MG is visualized on the UMAP embedding. Directions of the projected cell velocities on t-SNE are represented with arrows in the UMAP. D-E) Velocity UMAP representing the transcriptional velocity of *Lgals1* (D) or *St6gal1* (E) projected onto the 2D UMAP based on the dynamic model, with dark green indicating increased transcription and red indicating decreased transcription. The analysis was performed on 23,184 EpCAM⁺ cells from the MG of sorted C57BL/6J mice. F) Splicing dynamics across the different temporal stages experienced by the MG. G) Phase plot for the top 5 genes more represented in each MG lineage detected by the dynamical model. The function used from the scVelo package returns a structured table indexed by cell lineage, ranking genes based on similarity obtained during the dynamic model. H) Analysis of *LGALS1* expression across different epithelial and stromal mammary gland lineages.

**Figure S3:**
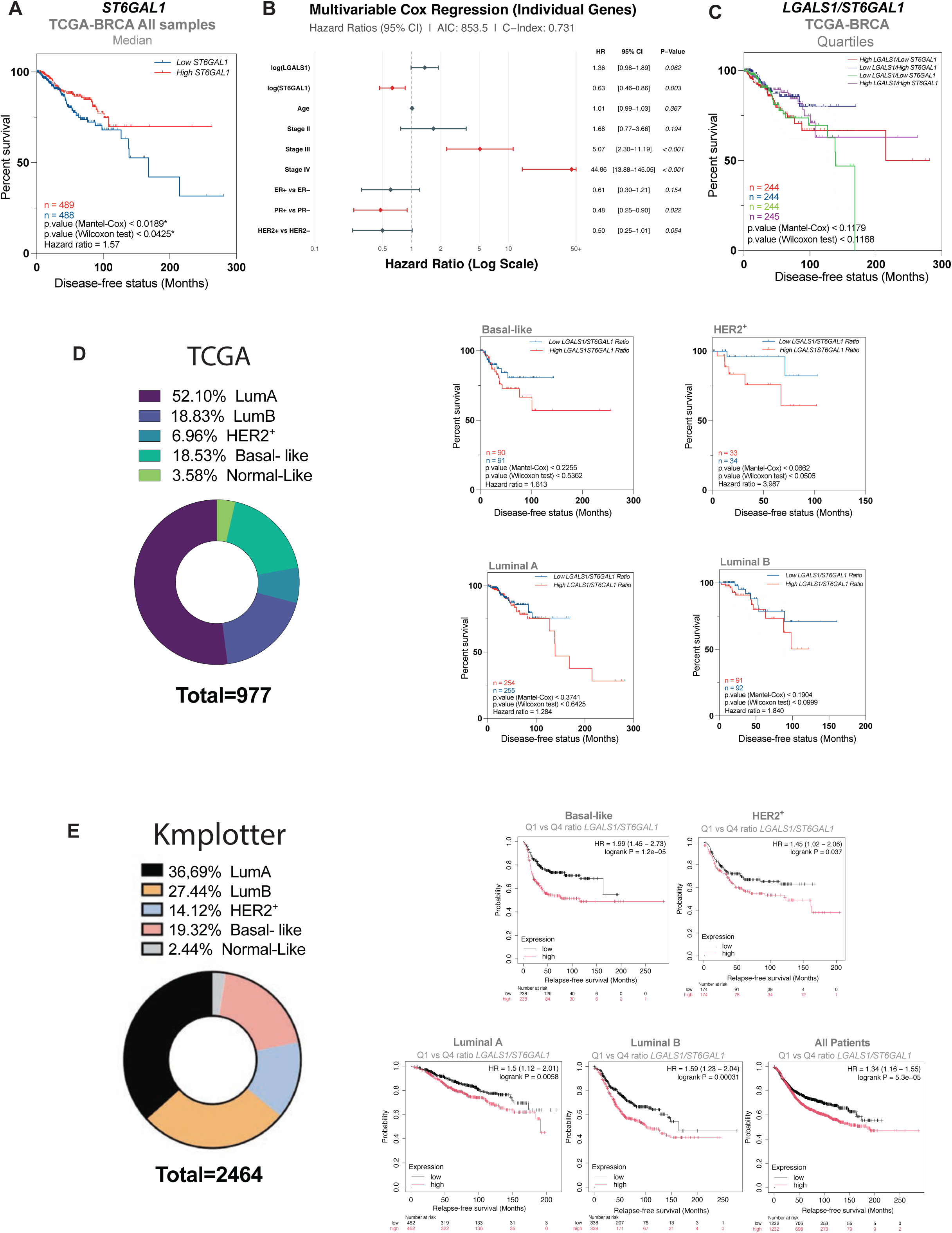
Prognostic value of *LGALS1/ST6GAL1* ratio after PAM50 stratification analysis. A) Kaplan-Meier DFS survival curve of patients with *ST6GAL1^high^* and *ST6GAL1^low^* expression (stratification of TCGA cohort: median *ST6GAL1* expression). B) Kaplan-Meier DFS survival curve of the four groups of patients stratified on median *LGALS1* expression, followed by median *ST6GAL1* expression. C) Kaplan-Meier DFS survival curve of patients with *LGALS1^high^/ST6GAL1*^low^ and *LGALS1^low^/ST6GAL1^high^* expression after PAM50 stratification of TCGA cohort (Median expression). D) Kaplan-Meier relapse-free survival (RFS) curve of patients with low and high *LGALS1/ST6GAL1* ratio (Q1 vs Q4) using the kmplotter dataset. **Statistical analysis:** (A, B, C and D) Disease-free survival was compared using the Mantel-Cox log-rank and Wilcoxon tests; hazard ratio (HR) was calculated from the survival curves.

**Figure S4:**
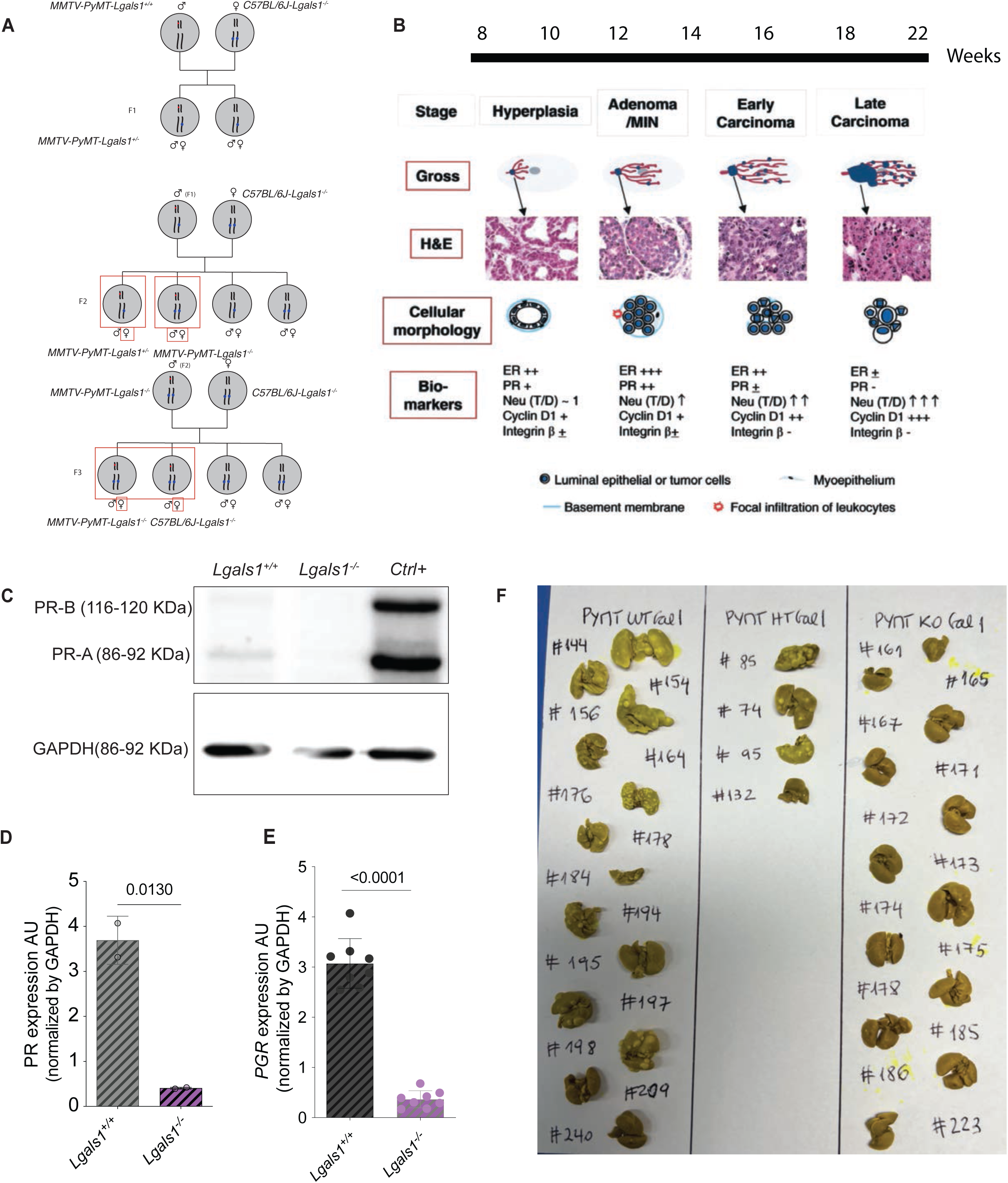
***MMTV-PyMT Lgals1^-/-^* mouse model development.** A) Schematic representation of the *PyMT-Lgals1*^-/-^ mouse model development strategy. B) Schematic representation of the different tumorigenic stages of MG and disease progression in the *MMTV-PyMT* mouse model. C) Western blot of PR A and B expression. D) Western blot analysis of PR A relative to GAPDH expression. E) qPCR analysis of PR mRNA expression F) Representative lung images after euthanasia, stained with picric acid to visualize macrometastatic foci in all genotypes. At end point all mice had tumors of similar size. **Statistical analysis:** (D-E) Unpaired Welch’s t-test. p-values < 0.05 are considered significant. Experiments were performed with n=2 (D) and n > 5 (E).

**Figure S5:**
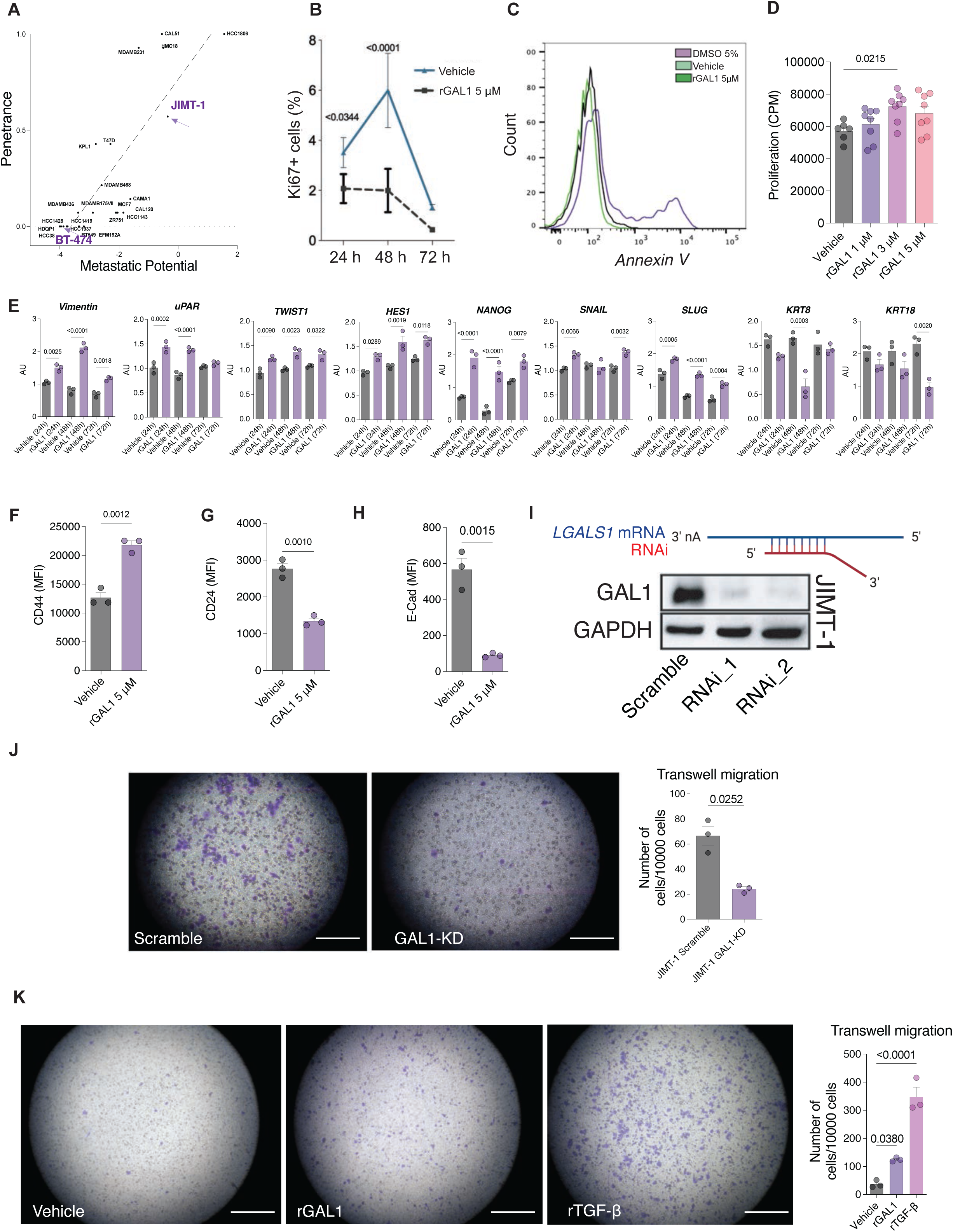
GAL1 triggers EMT and stemness in HER2^+^ human breast cancer cells. A) Plot of metastatic potential versus penetrance across different breast cancer cell lines. B) Percentage of Ki67^+^ cells (24, 48 and 72 h) after addition of 5 μM rGAL1 or vehicle. C) Frequency of annexin-V^+^ cells determined by flow cytometry. DMSO was used as positive control. D) Proliferation in BT-474 cells measured by ^3^ [H]-thymidine incorporation (72 h) after addition of 1, 3 and 5 μM rGAL1. E) qPCR of a list of EMT- and stemness-associated genes in JIMT-1 cells (24 h, 48 h and 72 h) treated with 5 μM rGAL1 or vehicle. F-H) Determination of CD44 (F), CD24 (G) and E-Cad (H) by flow cytometry (72 h) in JIMT-1 cells treated with 5 μM rGAL1 or vehicle. I) Western blot showing GAL1 expression in JIMT-1 cells following shRNA-mediated knockdown. J) Transwell migration of shRNA-silenced or scr JIMT-1 cells. K) Transwell migration of JIMT-1 treated with 5 μM rGAL1, vehicle (negative control) or TGF-ß_1_ (positive control). **Statistical analysis:** (C-G-H-J) Unpaired Welch’s t-test. (B-E-L) One-way ANOVA with Tukey’s post-test. p-values < 0.05 are considered significant. All experiments were performed with n ≥ 3 per group.

## METHODS

### Reagents and resources

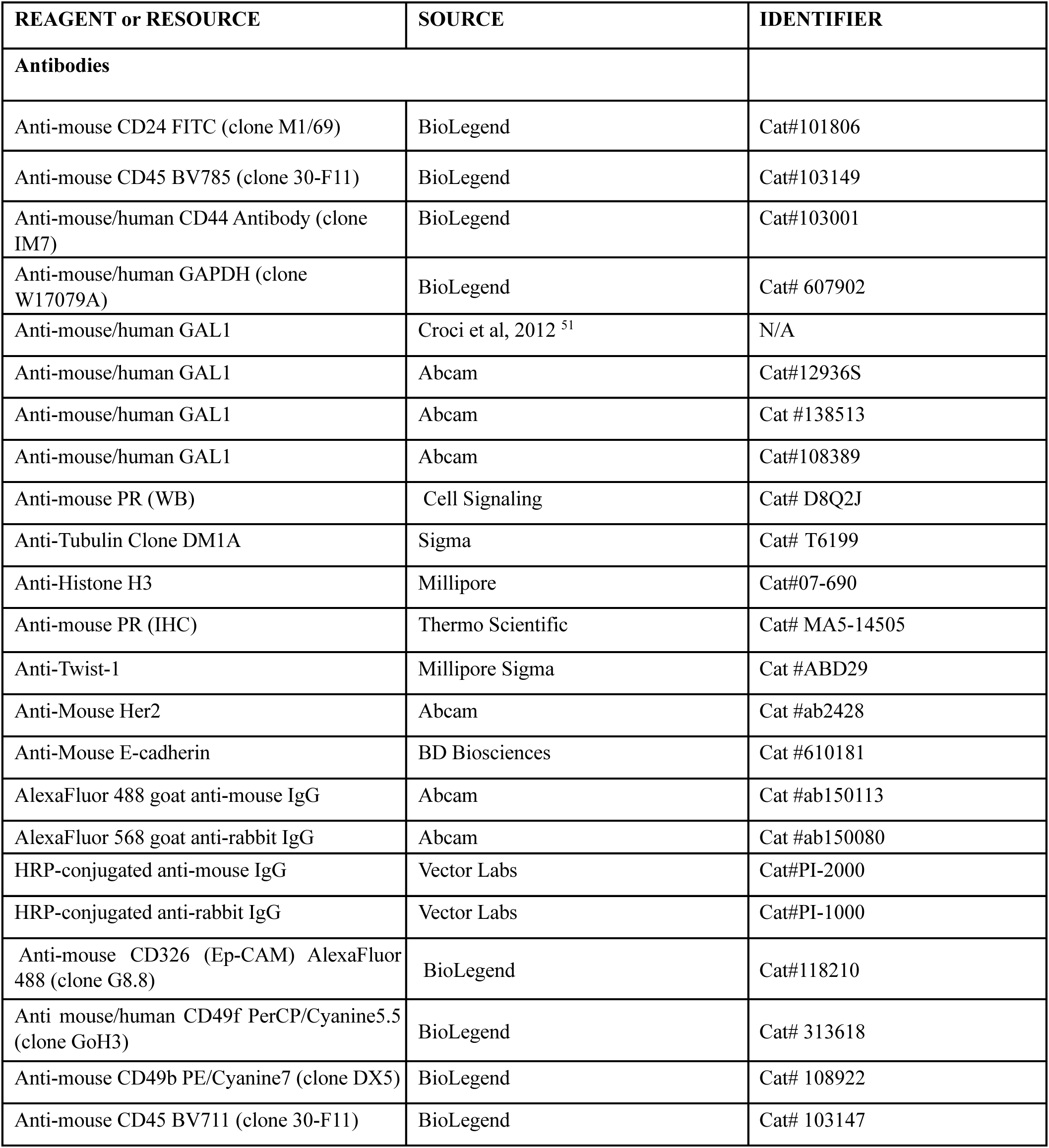
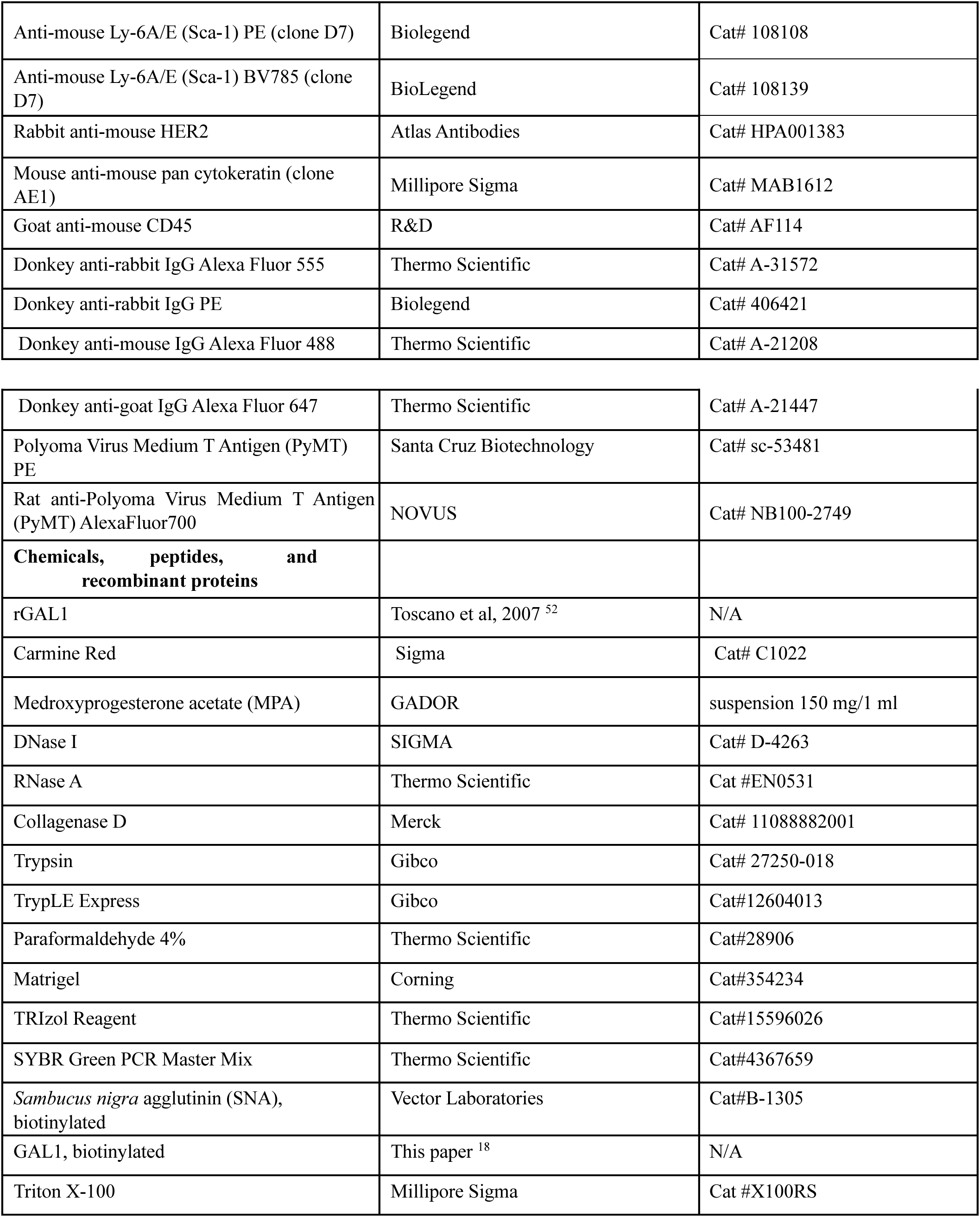
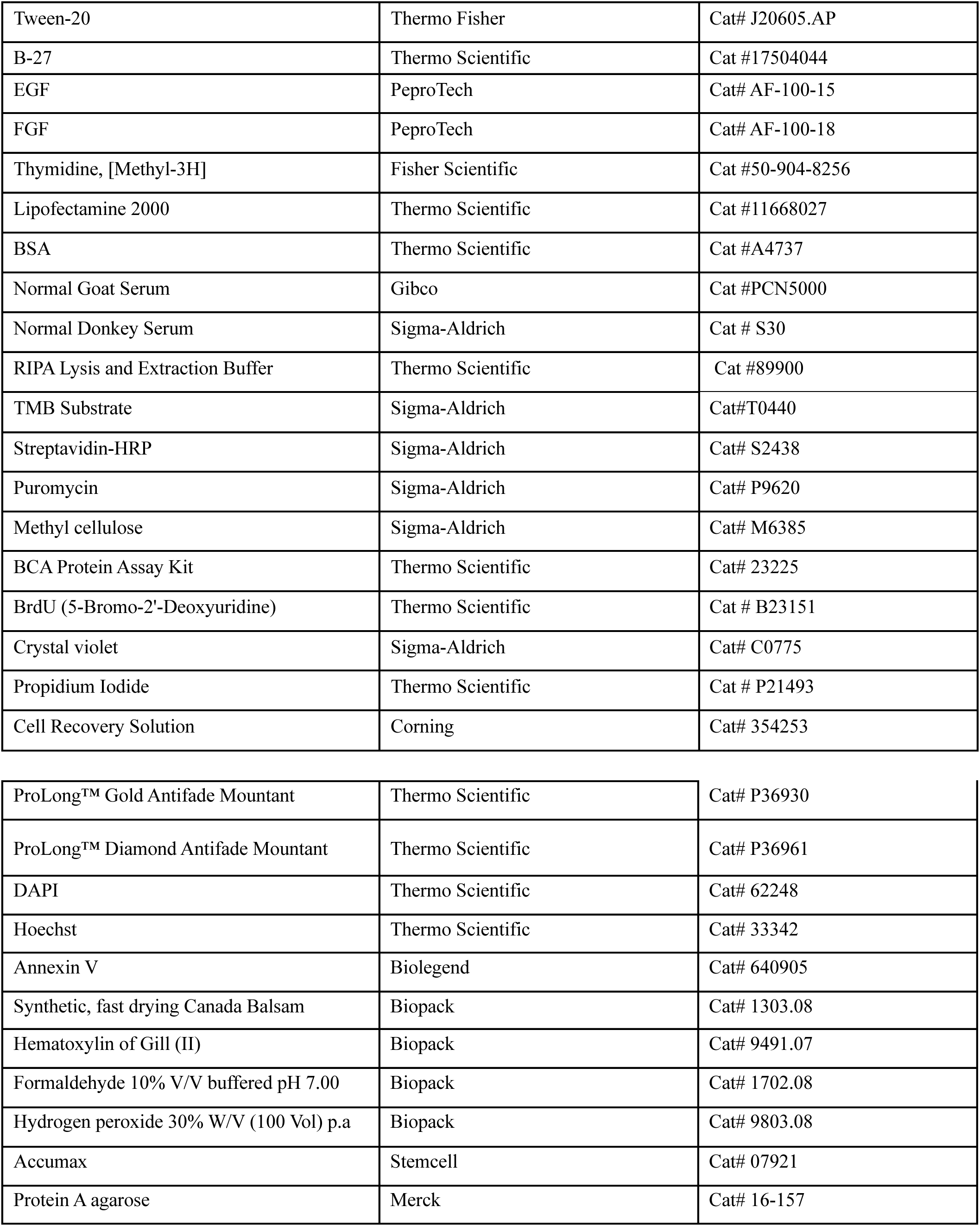
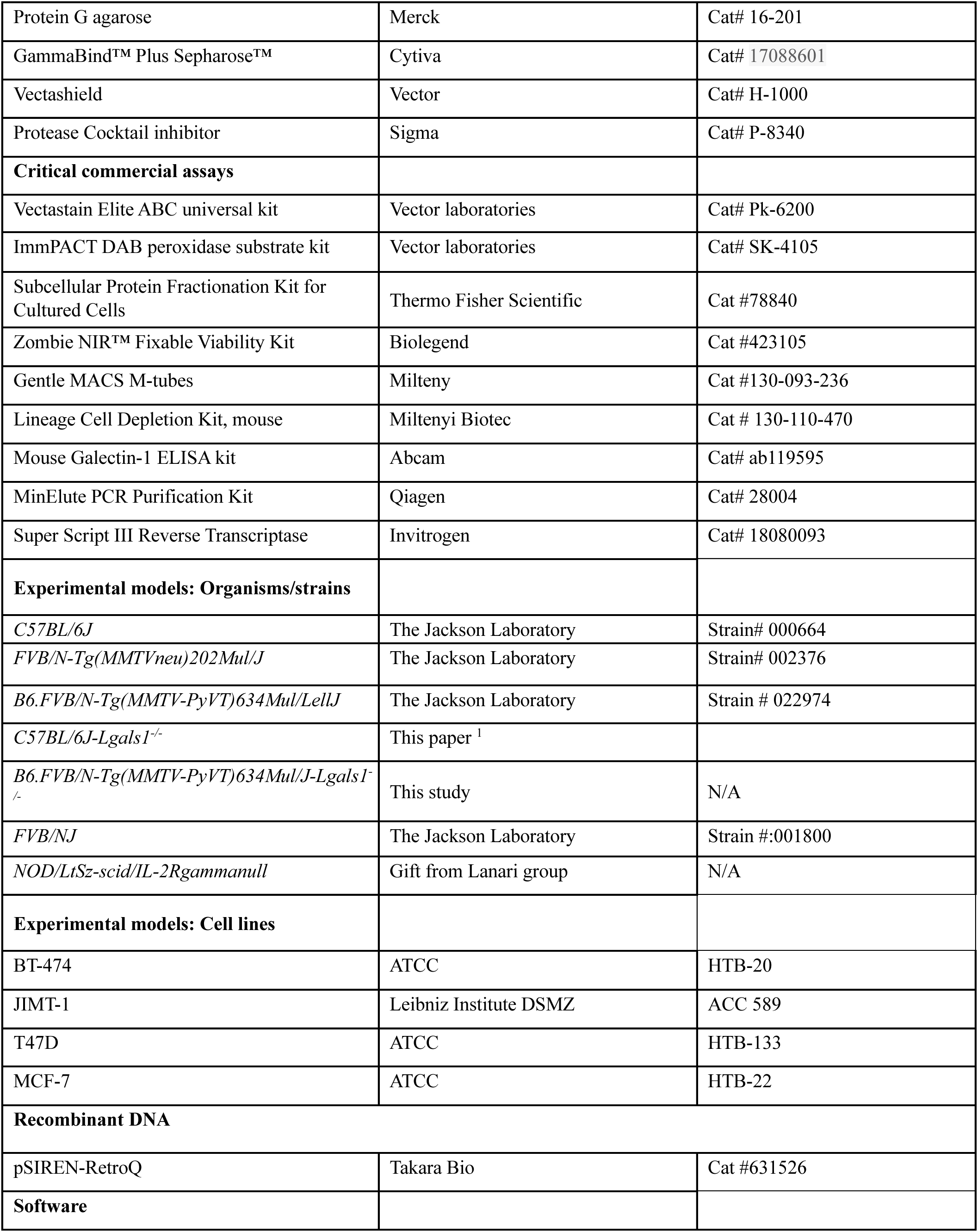
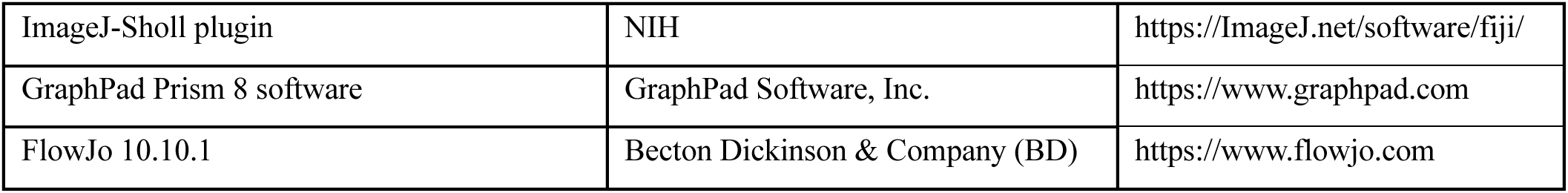

### Mouse models

B6.FVB-Tg(*MMTV-PyVT*) *634Mul/LellJ*-*Lgals1^-/^*^-^, was bred and crossed in our facilities for this study (Figure S4A). *MMTV-Neu* mice on FvB3 were bred and crossed at the Icahn School of Medicine at Mount Sinai and Albert Einstein College of Medicine. *NOD/LtSz-scid/IL-2Rgammanull* were donated by Claudia Lanari (IBYME-CONICET).

To obtain *B6.FVB-Tg(MMTV-PyVT)634Mul/LellJ/Lgals1^-/-^*transgenic mice we crossed the two parental strains (male: *B6.FVB-Tg(MMTV-PyVT)634Mul/LellJ (PyMT*) and female: *Lgals1^-/-^* C57BL/6 ^1^) (Figure S4A). In the F1 we obtained approximately 25% PyMT *Lgals1^+/-^* males (F1), which will be mated again with C57BL/6 *Lgals1^-/-^*females, to obtain 25% PyMT *Lgals1*^-/-^ litters (12.5% females). The resulting double transgenic PyMT *Lgals1^-/-^* females were compared with PyMT *Lgals1^+/+^*. Genotypes are confirmed in each litter by PCR, C57BL/6J *Lgals1*^+/+^ and *Lgals*1^-/-^ littermates are used as normal controls in each experiment.

Mice were maintained at specified pathogen-free (SPF) health status in individually ventilated cages at 21-22°C and 39-50% humidity. All animal procedures were approved by the Institutional Animal Care and Use Committee (IACUC) of the Icahn School of Medicine at Mount Sinai, the IBYME and Montefiore-Einstein. 8-week-old female mice were used to analyze the MG branching. For transgenic models, 14 to 18-week-old female mice were used as early (‘early lesion’; EL) stage mice and 24-week-old or older females with palpable tumors were used as late stage of cancer progression. Female littermates were randomly assigned to experimental groups. Tumors were not allowed to grow beyond the IACUC allowed limit of 1 cm^3^ per tumor.

### Cell culture

BT-474, T47D and MCF-7 cell lines were obtained from ATCC, while JIMT-1 cells were acquired from the Leibniz Institute DSMZ (German Collection of Microorganisms and Cell Cultures). JIMT-1, BT-474 and T47D were cultured in RPMI-1640 medium, while MCF-7 was cultured in DMEM-F12 (GIBCO), all supplemented with 10% fetal bovine serum (GIBCO). MCF-7 and T47D cell lines were also cultured with 1 μg/ml insulin (GIBCO). For passaging, cells were washed with PBS or physiological saline and incubated with TrypLE Express (GIBCO) for 4 min at 37°C. All cell lines were routinely tested for mycoplasma contamination using PCR-based assays.

### Hormonal induction of MG branching

Female *Lgals1^+/+^* and *Lgals1^-/-^* mice on both C57BL/6J were subjected to hormonal stimulation by subcutaneous injection of a suspension of medroxyprogesterone acetate (MPA; 15 mg) using a 25G needle in the right flank. Upon injection, MPA formed a subcutaneous depot that enabled slow and sustained release of the hormone. Age-matched non-injected mice were used as controls. After 21 days, animals were euthanized, and MG were harvested for flow cytometry or whole-mount analysis.

### Whole-mount staining of MG

Excised MG were carefully spread onto positively charged glass slides and fixed in Carnoy’s solution (60% absolute ethanol, 30% chloroform, 10% glacial acetic acid) for 4 h at room temperature. Fixed tissues were rehydrated through sequential incubation in 70%, 50% and 25% ethanol for 15, 5, and 5 min, respectively, followed by a 5-min wash in distilled water. Glands were stained overnight with carmine alum solution. After staining, samples were dehydrated through 70%, 95%, and 100% ethanol for 15 minutes each, cleared in xylene, and mounted in Canada balsam.

### TEB analysis

Terminal end buds (TEB), key hormone-responsive structures within the MG, were assessed manually using ImageJ software. Three randomly selected areas were analyzed per wholemount preparation, and the average number of TEB per area was calculated and used for subsequent graphical representation.

### Branching determination

Following whole mount staining of the MG, an image was acquired using a magnifying glass and a ruler to later adjust the scale. The image was opened using ImageJ (FIJI), transformed to 8-bit, and using the freehand tool, the lymph node was removed. Typically, the image had background noise that the plugin could easily confuse with MG branching. To avoid this, the background and any outliers were carefully subtracted while using the original image as a reference. Afterwards, the threshold value was adjusted until an adequate depiction of the gland was achieved. With the skeletonized version of the MG, an ending radius was determined by drawing a line from the start of the primary duct to the most distal point of the glandular epithelium. Because mammary branching is similar in morphology to neuronal ramifications, the Sholl Analysis for Neuroanatomy plugin was used to obtain the branching index.

### Flow cytometry and sorting of normal murine MG

To analyze the cellular lineages of the normal murine MG, we used 8-week-female C57BL/6J *Lgals1^+/+^* and *Lgals1^-/-^*, as reported by Ludwik et al ^53^. Before tissue manipulation, mice were grouped (mixing both *Lgals1^+/+^* and *Lgals1^-/-^* genotypes) for 24-48 h and synchronized by placing urine-saturated bedding from a male cage 24 h prior to the sacrifice. Vaginal cytology was performed on random mice to check that mice were in the same stage of their estrous cycle. Mammary glands were manually dissected using scissors following the removal of lymph nodes. Tissue dissociation was performed enzymatically in 7.5 mL of pre-warmed DMEM: F12 supplemented with 15 mg of collagenase D (2 mg/mL) and sterile penicillin/streptomycin (100x) for 2 h at 37°C. Following centrifugation (150 x g for 5 min), pelleted cells were resuspended in 1 mL of DMEM: F12 supplemented with DNase I and incubated for 5 minutes at 37°C. The resulting cell suspension was centrifuged (150 x g for 10 min) and washed once with PBS. Afterwards, the supernatant was aspirated, and the pellet resuspended in 1 mL of Accumax (Stem Cell). Samples were then divided equally into two 1.5 mL centrifuge tubes for incubation in a Shaking Incubator (Labnet VorTemp) at 37°C while shaking at 800 rpm for 10 min. Halfway through, large fibrous aggregates were removed. Cells were pelleted (500 x g for 5 min) and after separating the supernatants, samples were resuspended in a total volume of 0.5 mL TrypLE Express and incubated at 37°C for 3-5 min to complete isolation of mammary cells. The resulting cell suspension was filtered through a 70 μm cell strainer and centrifuged at 300 × g for 8 min. A total of 1 × 10⁶ cells per sample were plated in 96-well round-bottom plates (Greiner Bio-One) and stained with lineage-specific conjugated antibodies. Live cells were identified by exclusion of the viability dye (Zombie NIR -Biolegend). Cells were fixed with 4% paraformaldehyde and analyzed. Flow cytometric analyses were performed using either LSRFortessa (Becton Dickinson) or an Aurora Spectral flow cytometer (Cytek). Data acquisition was followed by analysis using FlowJo software (version 10.7.1; FlowJo, LLC). For cell sorting protocol, cell suspension was resuspended in PBS supplemented with 2% fetal bovine serum (FBS) and stained with the following antibody panel: anti-CD326 (Ep-CAM)-Alexa Fluor-488, anti-CD49f-PerCP-Cy5.5, anti-CD49b-PE-Cy7, anti-Sca-1-PE, and anti-CD45-BV711, along with Zombie NIR for cell viability. Fluorescence-activated cell sorting (FACS) was performed on a BD FACSAria™ cell sorter to isolate targeted populations. Post-sort, harvested cells were pelleted by centrifugation at 300 x g for 5 min and resuspended in TRIzol reagent for downstream purity analysis.

### RNAseq analysis of mouse MG RNA extraction

For RNA extraction, we used 8-to-12-week-old female previously grouped and synchronized C57BL/6J and *PyMT* mice, both *Lgals1^+/+^* and *Lgals1^-/-^*. First, mechanic tissue dissociation was performed in 1 ml TRIzol reagent in M-tubes gentleMACS (Miltenyi). After centrifugation (12,000 x g for 5 min at 4°C), the supernatant was collected and further processed according to the established protocol for TRIzol reagent. To evaluate the quality of the RNA extracted from our samples, we determined their RQN. Only samples with RQN > 7 were sent for scRNAseq to Macrogen (Korea).

### RNAseq analysis

Raw sequencing reads (FASTQ format) from cell lines were subjected to quality control and adapter trimming using Fastp ^54^. Transcript abundance quantification was performed using Kallisto ^55^ in pseudoalignment mode against the Ensembl mouse transcriptome reference (*Mus musculus*, GRCm38/mm10 assembly). Gene-level counts were aggregated from transcript-level estimates and imported into R for downstream analysis. Differential gene expression analysis was conducted using the Limma-voom pipeline ^56^, which applies variance modeling at the observational level to RNA-seq count data. For pathway enrichment analysis, gene set enrichment analysis (GSEA) ^57^ was performed with pre-ranked gene lists based on differential expression statistics. The Hallmarks gene set collection from the Molecular Signatures Database ^58^ (MSigDB) was used to identify enriched biological pathways.

### Chromatin immunoprecipitation

MCF-7 and T47D cells were processed for ChIP using 10 × 10⁶ cells per IP. Cells were crosslinked with 1% formaldehyde for 10-15 min at room temperature and quenched with 0.125 M glycine, followed by washes in cold PBS. Pelleted cells were lysed in soft lysis buffer (50 mM Tris-HCl pH 8.0, 2 mM EDTA, 0.1% NP-40, 10% glycerol, protease inhibitors) to isolate nuclei, which were then resuspended in nuclear lysis buffer (50 mM Tris-HCl pH 8.1, 10 mM EDTA, 1% SDS) and sonicated to obtain chromatin fragments of ∼200-500 bp. After centrifugation, samples were adjusted to 1mg/ml in dilution buffer (16.7 mM Tris-HCl pH 8.1, 167 mM NaCl, 1.2 mM EDTA, 1% Triton X-100, 0.01% SDS, protease inhibitors), precleared with Protein A/G agarose beads and salmon sperm (1 mg/ml) for 2 h, and 10% was reserved as input. Immunoprecipitation was performed overnight at 4 °C with 2.5 µg anti-GAL1 antibody (Abcam ab138513 or rabbit polyclonal ^51^) or rabbit IgG control, followed by incubation with Protein A/G beads. Beads were sequentially washed three times with low-salt buffer (20 mM Tris-HCl pH 8.1, 150 mM NaCl, 2 mM EDTA, 1% Triton X-100, 0.1% SDS), high-salt buffer (same composition with 500 mM NaCl), and LiCl buffer (10 mM Tris-HCl pH 8.1, 0.25 M LiCl, 1 mM EDTA, 1% NP-40, 1% deoxycholate). Chromatin was eluted in 0.1 M Na₂CO₃ and 1% SDS, and crosslinks were reversed overnight at 65 °C in the presence of 200 mM NaCl. Samples were treated with proteinase K (200 mM Tris-HCl pH 6.5, 50 mM EDTA) at 55 °C, and DNA was purified using the MinElute PCR Purification Kit (Qiagen) for downstream ChIP-qPCR analyses. Quantitative PCR (qPCR) analysis of ChIP DNA was performed using the QuantStudio real-time PCR system (Applied Biosystems). For each sample, two independent primer pairs were used to amplify regions within the *PGR* promoter A and two primer pairs targeting *PGR* promoter B. Reactions were set up in technical duplicates using appropriate SYBR Green master mix, following the manufacturer’s recommendations.

### Total extracts immunoprecipitation

MCF-7 cells were lysed in RIPA buffer supplemented with protease inhibitors. Cell lysates were precleared by centrifugation at maximum speed for 10 min at 4°C. Proteins were quantified in supernatants by the DC Protein Assay (Bio-Rad) and 1 mg of protein was incubated with 1 mg of primary anti-GAL1 antibody or irrelevant IgG overnight at 4°C with agitation. Following overnight incubation, 20 μl of effective GammaBind™ Plus Sepharose™ was added for 1 h and 30 min at 4°C with agitation. Afterwards, the beads were washed three times with PBS 0.1% NP40 to remove non-specifically bound proteins. Immunoprecipitated proteins were eluted from the beads by boiling in loading buffer for 5 min. Eluted proteins were resolved by SDS-PAGE using 14% polyacrylamide gels and analyzed by Western blot.

### Membrane and insoluble compartment immunoprecipitation

Cell extracts were prepared by homogenizing cells in 1% digitonin lysis buffer (25 mM Tris-HCl, pH 7.6, 150 mM NaCl, 1 mM EDTA, 1% digitonin), supplemented with protease inhibitors (0.3 μM aprotinin, 1 μM leupeptin, 1 μM pepstatin, 1 mM AESBF). These mild conditions allow the solubilization of membrane and cytosol and not cytoskeleton-associated structures and nuclei. Extracts were left on ice for 10 min and centrifuged at 14,000 x g for 10 min at 4°C. Supernatants constituted soluble cell extracts. Insoluble pellet corresponds to nuclei and cytoskeleton and was lysed with RIPA buffer supplemented with protease inhibitors. Proteins were immunoprecipitated as described above and immunocomplexes were analyzed by Western blot.

### immunofluorescence of GAL1 staining in cell lines

Cells were seeded onto 12 mm circular coverslips in 12-well plates and grown to 70-80% confluency. The culture medium was removed, and cells were washed twice with PBS prior to fixation. Cells were fixed using a solution of 4% (v/v) PFA and Triton X-100 (v/v) 0.2% for 10 min at room temperature, followed by permeabilization with 0.2% Triton X-100 in PBS for 10 minutes at room temperature. Nonspecific binding sites were blocked with 3% bovine serum albumin (BSA) in PBS for 1 hour at room temperature. Cells were then incubated overnight at 4 °C with anti-GAL1 primary antibody (Abcam Cat. ab108389; 1:100) diluted in incubation buffer (PBS + 1% BSA + 0.1% Triton X-100). After 18 h, coverslips were washed twice with PBS for 5 min each and incubated with the corresponding fluorophore-conjugated secondary antibody (Biolegend 1:400) for 1 h at room temperature in the dark. For nuclear staining, cells were incubated with Hoechst (1:500) for 10 min in the dark. Following two final washes with PBS, the coverslips were mounted onto glass slides using antifade mounting medium (Vectashield) and sealed. Images were obtained using an Olympus DSU fluorescent microscope.

### Analysis of public scRNA-seq datasets of mouse and human normal MG

To characterize *Lgals1* mRNA expression across epithelial and stromal cell populations in the normal MG, we reanalyzed publicly available single-cell RNA-seq datasets. Raw sequencing data from the murine MG studies by Bach *et al*. ^34^ (GSE106273) and Giraddi *et al*. ^26^ (GSE111113) were downloaded from the NCBI Sequence Read Archive. For the integrated murine epithelial atlas generated by Saeki *et a*l. ^33^ (GSE149949), the processed Seurat object was obtained from the UCSC Cell Browser. Because this atlas incorporates the Bach and Giraddi datasets, the cell annotations and UMAP coordinates provided in the processed object were retained for visualization and comparison across epithelial cell populations. The stromal compartment was examined using the processed count matrix and metadata obtained from Li *et al*. ^59^ (GSE150580). Human mammary epithelial populations were evaluated by reanalyzing the dataset from Nguyen *et al*. ^35^ (GSE113196). RNA velocity analysis of the Bach and Giraddi datasets was performed using Velocyto ^60^ to determine spliced and unspliced transcripts, followed by dynamic modeling with scVelo ^61^ . Cell-type annotations supplied by the original studies were used when available and evaluated based on established lineage markers. Gene expression and cluster-marker analyses were performed in Seurat, and *LGALS1*expression was compared across the annotated mammary cell populations.

### Immunofluorescence of normal murine MG

Excised mammary tissues were fixed in 4% (v/v) paraformaldehyde, embedded in paraffin, and sectioned for immunostaining. Sections were deparaffinized in xylene for 15 min, followed by rehydration through a graded ethanol series (100%, 90%, 80%, and 70%, 5 min each) and rinsed in distilled water. Antigen retrieval was performed in 10 mM citrate buffer, pH 6.0, by incubating the slides for 50 min at 90 °C in a water bath, followed by cooling at room temperature for 30 min. Permeabilization was carried out with Triton X-100 for 10 min at room temperature, followed by two 5-min washes (PBS + 0.02% Tween-20) under agitation. Tissue sections were then encircled using a hydrophobic barrier pen. Non-specific binding was blocked by incubating the sections for 60 min at room temperature in PBS containing 3% (w/v) BSA and 3% (v/v) normal goat serum. Primary antibodies, including anti-mouse α-SMA or rabbit polyclonal anti-mouse GAL1 were incubated overnight at 4 °C. The following day, sections were washed twice for 5 min in wash buffer with agitation, then incubated for 60 min at room temperature with the appropriate secondary antibodies diluted 1:1000 in blocking buffer. Then, slides were washed six times for 5 min in wash buffer. Finally, sections were mounted using ProLong Gold Antifade Reagent with DAPI (Invitrogen P36931) and covered with coverslips and allowed to dry at room temperature.

### Transcriptomic analysis

Transcriptomic data from breast cancer patients were downloaded from the TCGA-BRCA LEGACY repository using the R/Bioconductor package TCGAbiolinks ^62^. Data were filtered to remove duplicates, samples lacking disease-free survival (DFS) information, and male patients, resulting in a final cohort of 977 individuals. Differential gene expression analysis was performed using the Limma-voom package ^56,63^. Genes were considered up- or down-regulated if they exhibited a fold change (FC) greater than 2 and a false discovery rate (FDR) less than 0.001. GSEA of biological processes (GO terms) was conducted in a pre-ranked mode using the fast GSEA algorithm implemented in the fGSEA package ^57^. Pathways were deemed significantly modulated when FDR < 0.05. Heatmaps, bubble plots, and bar charts were generated using the ggplot2 package from Bioconductor. Patient survival analysis was also conducted using the online platform KMplotter (gene chip) (https://kmplot.com/analysis/)^64^ using the following parameters: **Affy id/ Gene symbol:** (Use ratio of two genes) *LGALS1* (201105_at)/*ST6GAL1* (201998_at); **Trichotomization**: Q1 vs Q4; **Survival**: RFS; **Follow up threshold:** All (Censored at threshold)**; Subtype - PAM50: e**ach separately.

The independent prognostic value of the *LGALS1/ST6GAL1* ratio was evaluated using multivariable Cox proportional hazards regression models, adjusting for age at diagnosis, tumor stage (I-IV), and receptor status (ER, PR, HER2). To validate the relevance of the ratio, we compared the goodness-offit of the ratio-based model against alternative models including individual gene expressions (*LGALS1* and *ST6GAL1*) using the Akaike Information Criterion (AIC) and the Concordance Index (C-index). Statistical significance was set at p < 0.05. All hazard ratios (HR) are reported with 95% confidence intervals (CI). To ensure a robust analysis, we excluded patients with missing data for key clinical covariates (Age, Stage, ER, PR, and HER2 status), resulting in a final cohort of 850 patients. HER2 status was determined by a hierarchical integration of FISH and IHC data to resolve equivocal cases where possible.

### PyMT protocols

#### Kaplan Meier survival curve

*B6.FVB-Tg(MMTV-PyVT)634Mul/LellJ Lgals^+/+^ (PyMT)* and *PyMT Lgals1^-/-^* were used. Tumor onset was monitored twice a week, and once the tumor appeared, the event was registered as 1 in the Kaplan Meier curve. Tumor-free mice were registered as 0. TFS (tumor-free survival %) was monitored until we reach 100% of events in 1. X-axis represents the time in weeks after birth for each mouse.

### Determination of micro and macrometastases in PyMT mice

PyMT *Lgals1^+/+^*and PyMT *Lgals1^-/-^* mice were euthanized at different times and MG were isolated for IHC and H&E staining, for whole mount protein and RNA extraction, and flow cytometry Lungs were excised, and PyMT^+^ CSC and DTC were stained for PyMT (Novus biologicals) EpCAM, CD24 and CD44 by flow cytometry

At endpoint (when tumor volume reached the ethical limit: 1 cm^3^), lungs were fixed and stained with Bowin Solution and macrometastases were determined using a magnifying glass. The number of foci were indicated per lung. N=15.

### Quantitative PCR

For qPCR, mRNA was extracted from cells by lysis and homogenization using TRIzol reagent (Thermo, Cat. No. 15596026), following the manufacturer’s instructions. DNA contamination was removed using DNase I. cDNA was synthesized using Super Script III Reverse Transcriptase (Invitrogen). For real-time PCR, the SYBR Green PCR Master Mix was used with an ABI System 7500 (Applied Biosystem). Gene expression was then normalized to reference genes, either GAPDH or Actin. The same protocol was applied for all qPCR performed during this study. Primers used are listed below:

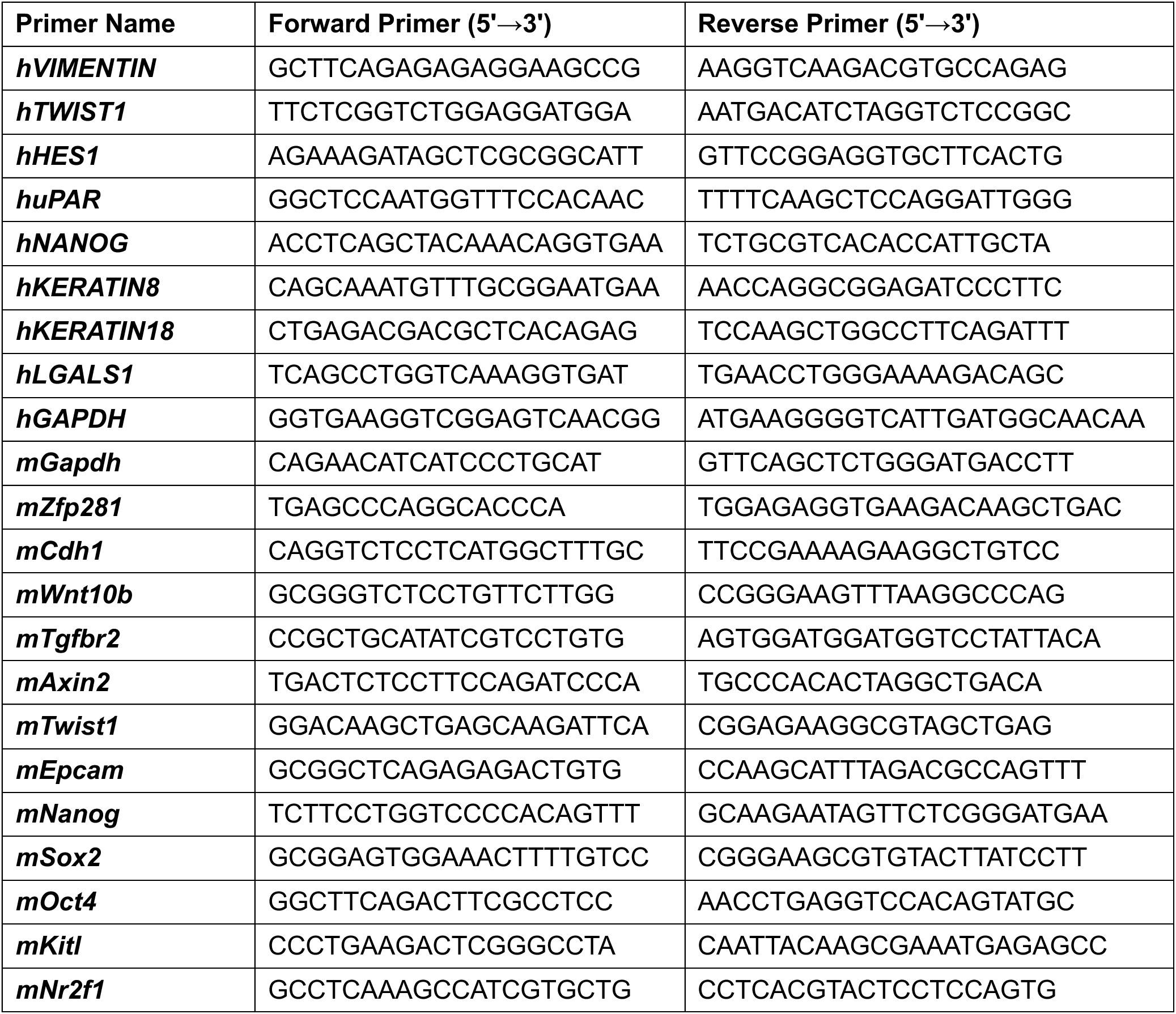
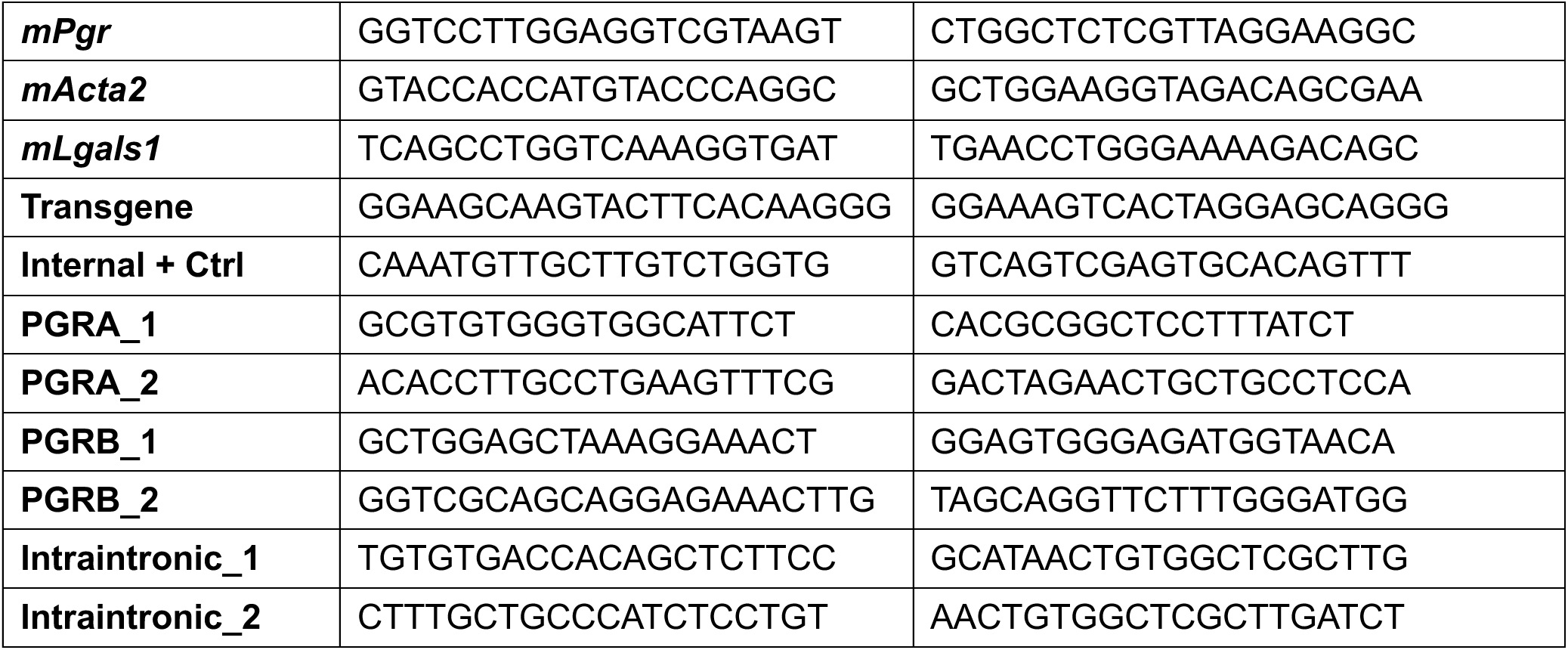

### Western blot

Cells were washed and then lysed in RIPA buffer (Thermofisher, PI89900) containing protease and phosphatase inhibitors (Thermofisher, 78440). Protein concentration was determined using a commercial MicroBCA assay kit with a BSA standard curve, following the manufacturer’s instructions (ThermoScientific # 23235). Next, proteins were resolved by SDS-PAGE, transferred to a nitrocellulose membrane (Cytiva Amersham Protran cat# 10600002), blocked with 5% bovine serum albumin (BSA) and incubated with primary antibodies. The following primary antibodies were used for immunoblotting or IHC: in house-generated anti-mouse/human GAL1 ^51^ (1:2000) or from obtained from Abcam; anti-GAPDH (Cell Signaling, 1:1000); anti-actin (Santa Cruz, 1:1000); anti-mouse PR (Cell Signaling); anti-tubulin (Sigma); Anti-histoneH3 (Millipore). The secondary antibodies used were HRP-conjugated anti-mouse (Vector Lab #PI-2000, 1:3000) and anti-rabbit (Vector Lab #PI-1000, 1:3000). Signal was detected using chemiluminescent substrate (ECL) (Thermo # 32106) and visualized with the GBOX imaging system (Syngene). Signal intensity was quantified using ImageJ software.

### PR immunohistochemistry

Histological sections were prepared from formalin-fixed, paraffin-embedded tissues. Sections from both *Lgals1^+/+^* and *Lgals1^-/-^* genotypes were deparaffinized in xylene (Stanton CAS 1330-20-7) overnight or for at least 2 h. Rehydration was performed by sequential immersion in 100%, 96% and 70% ethanol for 10 minutes each, using a Coplin jar. To block endogenous peroxidase activity, sections were incubated for 30 min in blocking solution (PBS or deionized water + 1% hydrogen peroxide), followed by two rinses in deionized water for 5 min each. Antigen retrieval was carried out in sodium citrate buffer (1 mM, pH 6.0) by placing the Coplin jar in a microwave on high power (800-watt) for 2-3 min, followed by a 2-3 min cooling period. This cycle was repeated three times. Afterward, slides were allowed to cool at room temperature for 20 min and then rinsed in deionized water for 10 min, followed by three 10-min washes in PBS 1x. Slides were then incubated for 30 min at room temperature in a humid chamber with blocking solution (PBS 1X + BSA 2.5%), followed by an overnight incubation at 4 °C in a humid chamber with the primary antibody (Invitrogen (Sp2) Cat# MA514505 anti-PR dilution 1:100 in PBS 1X). The next day, three 10-min washes in PBS were performed before incubation with the secondary biotinylated antibody (dilution 1:400 in PBS) for 60 min at room temperature. After another cycle of three 10-min washes in PBS 1x, slides are incubated with avidin-biotin complex (ABC method) for no more than 30 min at room temperature. After a final cycle of three 10-min washes in PBS, slides were incubated with DAB reagent (Sigma-Aldrich) until the desired staining intensity was achieved, then rinsed with water to stop the reaction. Finally, slides were counterstained with hematoxylin of Gill (II) for 30 sec, dehydrated, and mounted with synthetic, fast-drying Canada Balsam.

### Glycophenotyping

Glycophenotyping was performed as previously described ^65^. Cell lines were incubated with PBS-EDTA 15 mM to promote detachment from the plate without disrupting potential glycosylated membrane proteins. They were seeded in a 96-well multiwell plate and centrifuged at 300 x g for 8 min. Next, 100,000 cells were incubated for 30 min at 4°C with the following biotinylated lectins: *Sambucus Nigra* agglutinin (SNA, 10 μg/ml, Vector # B-1305-2) and rGAL1 (GAL1, 40 μg/ml. Subsequently, binding of each lectin was detected by flow cytometry. The relative median fluorescence intensity (rMFI) was calculated using the following formula: (median intensity of the cells of interest / median intensity of the negative control). To evaluate glycan-dependent GAL1 binding, lactose (30 mM) was used as a competitive inhibitor.

### Cell proliferation

To assess cell proliferation, 15,000 cells were seeded per well in octuplicate into 96-well plates. On the following day, cells were treated with recombinant GAL1 (rGAL1) at concentrations of 1, 3, and 5 µM or with vehicle control for 72 h, with a final volume of 150-200 µl per well. After treatment, 100 µl of culture medium were removed and replaced with 100 µl of fresh medium containing 1 µCi of tritiated thymidine ([³H]-thymidine) (Fisher Scientific #50-904-8256). Cells were subsequently harvested, and radioactivity was measured using a 1414 Liquid Scintillation Counter (PerkinElmer).

### Cell cycle analysis

To analyze the cell cycle, BrdU (Thermo # B23151) and propidium iodide (PI) (Thermo #P21493) staining was performed following this protocol. Cells grown under optimal conditions were incubated with BrdU at a final concentration of 30 μM for 30 to 60 min. As BrdU is light-sensitive, it was added in the dark, and all subsequent incubations were conducted under low-light conditions to avoid photodamage. The pulse time and BrdU concentration can vary depending on cell type and doubling time, typically ranging from 15 min to 2 h and from 10 μM to 100 μM. Following BrdU incorporation, the BrdU-containing medium was removed, and the cells were washed once with PBS. Cells were then trypsinized and collected. For permeabilization, the cell pellet was resuspended in 0.3 ml PBS, followed by the slow addition of 0.7 ml of cold 100% ethanol while gently mixing. After centrifugation at 4,000 rpm for 2 min, the supernatant was removed. Cells were then incubated in 0.5 ml of 2N HCl containing 0.5% Triton X-100 for 30 min at room temperature. After centrifugation and removal of the supernatant, cells were resuspended in 0.5 ml of 0.1 M sodium tetraborate for 2 min and centrifuged again. Cells were washed once with PBS containing 1% BSA, resuspended in PBS with 1% BSA and 0.5% Tween-20, and incubated with anti-BrdU mAb for 1 h at room temperature. After another wash with PBS 1% BSA, cells were incubated for 30 min with the secondary antibody. Finally, cells were resuspended in 0.5 ml PBS containing 10 μg/ml RNase A (Thermo # EN0531) and 20 μg/ml propidium iodide. Samples were incubated in the dark at room temperature for 30 min before acquisition by flow cytometry or stored at 4 °C until analysis.

### Cancer stem cell determination

Tumor cells were detached from the culture plate using TrypLE Express (GIBCO #12604013) for 5 min in a humidified incubator. After detachment, cytometry buffer using the following mAb: anti-CD24 (Clone M1/69 - BioLegend) at a 1:200 dilution, anti-CD44 (Clone IM7 - BioLegend) at 1:200 and analyzed by flow cytometry.

### Mammosphere formation assay

The capacity of JIMT-1 cells to form three-dimensional tumor spheres was evaluated following 72-h treatment with 5 µM rGal1. Between 0.5 and 1 × 10³ JIMT-1 cells were seeded in DMEM/F12 supplemented with B-27 (Thermo #17504044), EGF (20 ng/ml), FGF (20 ng/ml) (Peprotech) without serum. Cells were seeded in 24-well plates using 0.5 ml of medium per well. Tumor sphere growth was monitored over time, with daily gentle agitation of the plate to prevent adherence to the substrate or between spheres. After 1-2 weeks, the mammospheres were photographed and quantified under a microscope, considering both their number and size.

### Migration assay

Transwell migration assays were performed using 8 μm pore inserts (Millipore #PSET010R1) placed in 24-well plates. Tumor cells were seeded in the upper chamber of the transwell insert using RPMI medium containing 0-1% FBS (Gibco # 16000044). Between 3 × 10⁴ and 5 × 10⁴ cells were seeded in 200 μl of culture medium. To induce migration, a serum gradient was created by placing 0.5 ml of RPMI (Gibco # 61870036) supplemented with 20% FBS in the lower chamber. Cells were incubated overnight in a humidified incubator. After 24 h, transwells were removed, and residual medium was discarded. Cells were fixed and stained with 0.05% crystal violet (Sigma #C0775) in 20% methanol for 10 min at room temperature. Cells remaining in the upper chamber were carefully removed using a cotton swab. Migrated cells on the underside of the transwell membrane were visualized, photographed under a microscope, and counted.

### Scratch wound migration assay

JIMT-1 cells were seeded at an appropriate density (around 10-20K cells per well) on an Incucyte® Imagelock 96-Well Plate (Sartorius, BA-04855) in a standard cell incubator for between 6-18 h. Then, using the Incucyte® 96-Well Woundmaker Tool the wound was created in every well. After wounding, containing rGAL1 was incorporated. The plate was placed into the Incucyte® Live-Cell Analysis System, and a repeat scanning was scheduled (every 2 h for 24-48 h). Images were analyzed using the live-cell analysis software and graphs were obtained.

### GAL1 knockdown

GAL1 knocked down (GAL1-KD) cells were generated using a short hairpin RNA (shRNA) specific to GAL1, while control cells were transfected with a scrambled oligo, both cloned into the pSIRENRetroQ vector (TakaraBio #631526). Cells were transfected using Lipofectamine 2000 (Thermo #11668027) in serum-free Optimem (Thermo #31985062) medium for 24-48 h. Following transfection, selection was performed using puromycin (5 µg/ml), and clones exhibiting the lowest GAL1 expression were isolated by limiting dilution. GAL1 expression in KD clones, were assessed by qPCR and Western blot.

### Experimental metastasis

Breast cancer cells were cultured in RPMI (Gibco # 61870036) with 10% FBS and resuspended in PBS or serum-free medium at 1×10^6^ per 100 µl. *NOD/LtSz-scid/IL-2Rgamma^null^* mice (8-10 weeks old) were anesthetized with a mixture of ketamine (100 mg/kg) and xylazine (10 mg/kg) administered intraperitoneally. Cells were inoculated via tail vein injection and monitored until recovery. At the designated time point (6 weeks), mice were euthanized, and lungs were collected for analysis. The number and nature of macro-metastases were evaluated by staining with Bowin solution.

### Transgenic *MMTV-Neu* mouse model

Female *MMTV/Neu* transgenic mice aged 14-18 weeks were used to represent early-stage (“premalignant”, EL) lesions, while animals aged 20 weeks or older were considered to be at late-stage cancer progression (PT) ^1^. Mice were euthanized using isoflurane anesthesia followed by cervical dislocation. All five MG pairs were examined for the presence of small visible lesions or palpable tumors. Mice were perfused with PBS, and organs were collected. For histopathological analysis, tissues were fixed in 4% (v/v) PFA (Thermo Scientific #28906) for 24 h, processed, embedded in paraffin, and sectioned. For FACS and primary cell culture preparations, complete MG, primary tumors, and/or lungs were digested in 0.15% collagenase IA (SIGMA, C-9891) with 2.5% bovine serum albumin (BSA) at 37°C under agitation for 30 min. Red blood cells were lysed using RBC lysis buffer (Thermo # 00-4333-57) for 2-5 min, and the resulting cell suspensions were filtered through a 40 µm strainer, passed through a 25-gauge needle, and counted using a Neubauer chamber.

### RNA-seq analysis of EL and PT organoids

For transcriptomic profiling, we used processed RNA-seq data generated in Nobre et al. ^39^. To investigate gene expression programs activated in early lesions (EL) versus primary tumors (PT) in the *MMTV-Neu* model, bulk RNA sequencing was performed on mammospheres derived from EL and tumorspheres derived from PT, which recapitulate the *in vivo* behavior of EL and PT, respectively. Differential expression analysis identified 4,290 significantly regulated genes (adjusted p value < 0.05 and fold change > 2 or < 0.5), including 2,873 upregulated and 1,417 downregulated genes in EL compared to PT spheres.

### Three-dimensional (3D) culture and immunofluorescence

*MMTV-Neu* mice (EL - 14-18 weeks old or PT - >20 weeks) were sacrificed using CO₂, and mammary epithelial cells (MEC) were isolated as previously described ^39^. A total of 5.0 × 10⁴ MEC were seeded in 400 μl assay medium (DMEM/F12 (Gibco #21041025) supplemented with 5% horse serum (Gibco # 26050088), 1% penicillin-streptomycin (Gibco # 10378016), 20 ng/ml EGF (Peprotech), and 2% Matrigel (Corning # CLS356234) on 8-well chamber slides precoated with 40 μl Matrigel. After 24 h, organoids were treated with rGAL1 at concentrations of 1, 3, 5, or 10 μM or vehicle control every 48 h for one week. For immunofluorescence staining of 3D organoids and mammary glands, samples were fixed with 4% (v/v) PFA for 20 min at room temperature in the presence of phosphatase and protease inhibitors and then permeabilized using 0.1% Triton X-100 (Millipore #X100RS) in PBS for 20 min. Blocking was performed for 1 h in PBS-based immunofluorescence wash buffer (130 mM NaCl, 7 mM Na₂HPO₄, 3.5 mM NaH₂PO₄, 7.7 mM NaN₃, 0.1% BSA, 0.2% Triton X-100, 0.05% Tween-20) containing 10% normal goat serum (Gibco, PCN5000). Primary antibodies included:

E-cad (BD Biosciences, 610181; 1:100), TWIST1 (Millipore, ABD29; 1:100), in-house generated GAL1 (1:300), and HER2 (Abcam, ab2428; 1:100). Secondary antibodies used were Alexa Fluor 488 goat anti-mouse (Abcam ab150113; 1:1000) and Alexa Fluor 568 goat anti-rabbit (Abcam ab150080; 1:1000). Chambers were removed and samples were mounted using ProLong Gold Antifade reagent with DAPI. 3D organoids were imaged using a Leica SP5 confocal microscope with Leica software.

### Differential interference contrast (DIC) microscopy

Time-lapse imaging of 3D organoids treated with rGAL1 or vehicle control was performed over a 24-h period using a Zeiss LSM 880 confocal microscope equipped with Airyscan and a 10X/0.45 objective lens with 4X zoom. Four positions per condition were recorded in parallel, capturing images every 20 min, adjusting for cellular depth. Environmental conditions were maintained at 37 °C with 5% CO₂. Image acquisition and export (as uncompressed AVI files) were conducted using Zen 2.1 software. Videos were generated using Zen 2.1 or ImageJ. Imaging was performed at the Microscopy Facility of the Icahn School of Medicine at Mount Sinai (New York, NY, USA).

### GAL1 neutralization in the *MMTV-Neu* model

MMTV-Neu mice (EL - 14-18 weeks old females) were treated twice a week i.p. with anti-GAL1 mAb ^13,14^ or IgG2b isotype control (BioXCell Cat# BE0086) for 4 weeks (dose per mice: 30 mg/kg). At the end point of the experiment, mice were intracardially bled to collect blood for circulating tumor cells (CTC) isolation and subsequent plasma for analysis. GAL1 detection was performed in plasma using a GAL1 ELISA kit (Abcam Cat# ab119595) to confirm reduction of circulating GAL1 in mice treated with the anti-GAL1 mAb. For CTC isolation, the collected blood was treated with RBC lysis buffer to remove RBCs for 2-5 min on ice and neutralized with MACS buffer (PBS with 0.5% BSA and 2 mM EDTA). The remaining cells were counted, and immune lineage cells were removed using a MACS lineage depletion kit (Miltenyi Biotec Cat# 130-110-470). The unstained, negative fraction was cytospined (500 rpm for 10 min) onto microscope slides and fixed with 4% (v/v) PFA at room temperature for 10 min. The slides were washed with PBS and non-specific binding was blocked by incubating slides for 60 min at room temperature in PBS-Tween (PBS-T) containing 5% BSA (Fisher Scientific, Cat# BNP1605-100), 3% donkey serum (Sigma-Aldrich Cat# S30), and 0.05% (v/v) Tween-20 (Thermo Fisher, Cat# J20605.AP). Primary antibodies (anti-mouse HER2 1:200 (Atlas Antibodies, Cat. No. HPA001383), anti-mouse pan-cytokeratin 1:100 (Sigma Aldrich, Cat# MAB1612), and anti-mouse CD45 1:500 (R&D, Cat. No. AF114) were applied and incubated overnight 4 °C. After 24 h, slides were washed three times for 5 minu with PBS-T with agitation, then incubated for 2 h at room temperature with the appropriate secondary antibodies diluted 1:1000 in blocking buffer. Afterwards the slides were washed three times for 15 min each in wash buffer with agitation. Slides were then stained with DAPI (Thermo Fisher Cat# 62248) diluted 1:1000 in PBS for 5 min at room temperature, washed with PBS, mounted with ProLong Diamond Antifade (Invitrogen, P36961), covered with a coverslip and allowed to dry at room temperature.

### Statistical analysis

For comparisons between two groups, a Student’s *t*-test and Mann-Whitney *U*-test. For experiments involving three or more groups, one-way analysis of variance (ANOVA) followed by post hoc multiple comparisons with Dunnett’s test (comparing all groups to a control group) or Tukey’s test (comparing all groups to each other) was applied. Unless stated otherwise, experiments were performed in triplicate (n ≥ 3). Statistical analysis was performed using GraphPad Prism 10 software. Kaplan-Meier curves were generated in GraphPad Prism 9 and analyzed using Mantel-Cox (log-rank) and Wilcoxon tests.

